# QuantiTrack: A unified software to study protein dynamics in living cells

**DOI:** 10.64898/2026.02.19.706877

**Authors:** David A Ball, Kaustubh Wagh, Diana A Stavreva, Le Hoang, R Louis Schiltz, Raj Chari, Razi Raziuddin, Davide Mazza, Arpita Upadhyaya, Gordon L Hager, Tatiana S Karpova

## Abstract

Linking the spatiotemporal dynamics of proteins in live cells to biological function is a fundamental challenge in biology. Single molecule tracking (SMT) has emerged as a powerful technique to investigate protein dynamics at the single molecule level. However, SMT analysis often requires expertise in biophysical modeling and programming, and integrating results from different analyses can be challenging. To address these barriers, we developed QuantiTrack: a MATLAB-based SMT analysis software with a simple graphical user interface. This provides a much-needed end-to-end solution where a user can load a movie, detect and track single molecules, and perform complementary downstream analyses within a standardized workflow. QuantiTrack includes quantitative metrics for selecting detection and tracking parameters and troubleshooting experimental design, and includes a detailed step-by-step User Guide. We used simulations to demonstrate how signal intensity, labeling density, and motion blur affect detection and tracking fidelity. Using multi-state simulations, we further benchmarked complementary methods to identify distinct mobility states from heterogeneous trajectory populations. Finally, we applied QuantiTrack to real experimental data where we address how the glucocorticoid receptor (GR), a hormone-regulated transcription factor, responds to treatment and washout of its cognate hormone. Hormone washout results in rapid (in minutes) downregulation of GR target genes to basal levels. By integrating complementary analyses within QuantiTrack, we showed that hormone washout substantially reduced the bound fraction of GR, its occupancy in the mobility state associated with GR activation, and dwell times. Together, these analyses showcase QuantiTrack as an integrated platform for extracting biologically meaningful measurements from single molecule trajectories.

## INTRODUCTION

The living cell contains billions of proteins that are constantly moving, interacting with other proteins, nucleic acids, and small molecules to maintain cellular homeostasis and help the cell respond to external stimuli. Today, single molecule tracking (SMT) has become the state-of-the-art technology to study intranuclear protein dynamics across species ranging from yeast^1^, drosophila^2^, and zebrafish^3^, to mammals^4^ because it provides a direct readout of the motion of individual molecules.

For the same set of SMT data, different analytical approaches allow one to extract distinct, complementary information that quantifies the spatiotemporal dynamics of their protein of interest. These include measures of spatial mobility such as mean-squared displacements (MSD) or diffusivity distributions^5–14^, the fraction of bound molecules^5,15^, the anisotropy in turning angles^16,17^, and binding lifetimes^18,19^ (‘dwell’ times). Specifically, those approaches characterize the speed and geometry of the search for target sites, dynamics of free, confined, and bound molecules, and the transitions between bound and diffusive states. The current goal is to connect these parameters to the physiological role of the protein of interest. The success of this endeavor relies on proper data collection and rigorous analysis and interpretation.

Several groups have developed complementary analysis tools that are often used in conjunction with one another. To do this, one must run the same data through a roster of different programs, which creates a bottleneck in the SMT workflow. First, this requires a sophisticated understanding of programming languages along with various biophysical models, something that is not always readily available within a molecular biology laboratory. Second, it slows down the quantification, as there is no easy way to transfer the data between programs, and many steps must be repeated in different programs. Finally, comparing similar parameters obtained from different programs and maintaining uniformity therein is challenging.

To overcome this, we developed a comprehensive analysis software called QuantiTrack. This is a MATLAB-based software that runs through a graphical user interface (GUI), where users can identify and track single molecules from a movie, perform quality control (QC), and analyze tens of thousands of single molecule trajectories across hundreds of cells with a few clicks and minimal knowledge of MATLAB. QuantiTrack not only allows users the freedom to choose between multiple tracking algorithms but also to use well defined QC metrics to systematically determine the detection and tracking parameters. The data are stored in a convenient table format, which serves as the input to downstream analyses. Within QuantiTrack, we have bundled the following features that we believe are essential to quantify the spatiotemporal dynamics of a protein-of-interest:

1. Kinetic modeling and Spot-On to calculate fractions of bound and diffusive molecules
2. Iteratively fitting the step size distribution (or van Hove correlation) to estimate the distribution of MSDs using the Richardson-Lucy algorithm (RL analysis)
3. Classifying trajectories into distinct mobility states using perturbation expectation-maximization version 2 (pEMv2)
4. Quantifying the anisotropy in the distributions of turning angles over time and space
5. Fitting ensemble MSD to anomalous diffusion models
6. Measurement of photobleaching rates
7. Calculating photobleaching-corrected dwell time distributions

QuantiTrack comes with a detailed User Guide and simple interface, which makes it highly accessible to every molecular biologist, and generates publication quality figures and movies.

We provide a systematic framework for selecting detection and tracking parameters and evaluating the associated tradeoffs. Using simulated SMT timelapse images with known ground truth trajectories, we determined how signal intensity, motion blur, diffusion coefficient, and labeling density affect detection and tracking fidelity. We further used multi-state simulations to compare the complementary analyses implemented in QuantiTrack and evaluate the strengths and limitations of each approach.

As a practical example, we used QuantiTrack to investigate the dynamics of the glucocorticoid receptor (GR, a steroid hormone receptor). GR is a ligand-regulated transcription factor (TF) that binds specific DNA sequence motifs within enhancers to regulate target genes. We have previously demonstrated that the timing and duration of glucocorticoid delivery is an important and often overlooked aspect of treatment protocols^20–22^. Many GR target genes are rapidly activated within minutes of hormone treatment and return to basal levels shortly after hormone washout^20,21,23^. Using FRAP, we showed that hormone washout results in faster fluorescence recovery indicating shorter GR binding lifetimes (referred to as ‘dwell times’ in this manuscript), and reduced GR binding via chromatin immunoprecipitation (ChIP)^20^. In this study, we validated these findings using SMT, and showed that hormone washout results in a substantial reduction in the GR bound fraction, binding lifetimes, and GR occupancy in the lowest mobility state which correlates with the active form of TFs^7,24,25^. Together, these findings show that a unified SMT analysis platform such as QuantiTrack enables systematic investigation of protein dynamics in cells and clarifies how dynamic changes relate to function, making SMT accessible to a broad community of biologists.

## RESULTS

### QuantiTrack: A GUI-based SMT analysis platform

SMT experiments are typically performed by confining the excitation light to a thin sheet (∼micrometers for Highly Inclined Laminated Optical [HILO] Sheet microscopy, ∼100 nm for Total Internal Reflection Fluorescence [TIRF] microscopy) around the focal plane to reduce out-of-plane fluorescence and achieve high signal-to-noise ratios (SNR) **(Figure 1a)**. The initial output of an SMT experiment is shown in **Figure 1b** where single molecule trajectories are represented as kymograms, showing stable trajectories with long residence times along with transient and highly mobile trajectories. Stopped molecules appear as long stripes while faster diffusing molecules appear as short squiggles **(Figure 1b, Movie 1)**. The typical workflow for SMT analysis involves using a software such as u-Track^26^ or TrackMate^27^ to identify single molecules within the movie and to connect them together into tracks **(Supplementary Figure 1)**. Multiple software programs are then used to analyze various spatiotemporal characteristics of these trajectories, including but not limited to, dwell time distributions, turning angle anisotropy, bound fractions, other measures of diffusivity distributions (a non-exhaustive list of software can be found here^28^). Individual research groups must often write custom scripts to perform quality control and to generate publication-worthy figures and movies. The required programming skills are not readily available in every molecular biology laboratory, creating a high barrier-to-entry for a molecular biologist looking to perform SMT. Here, we combine all these features into a GUI-based tool where a user with minimal programming knowledge can perform all these steps with relative ease and obtain quantitative information about their biological system of interest. We call this software QuantiTrack **(Figure 1c)** and it is based on the original framework of TrackRecord^15^.

**Figure 1:**
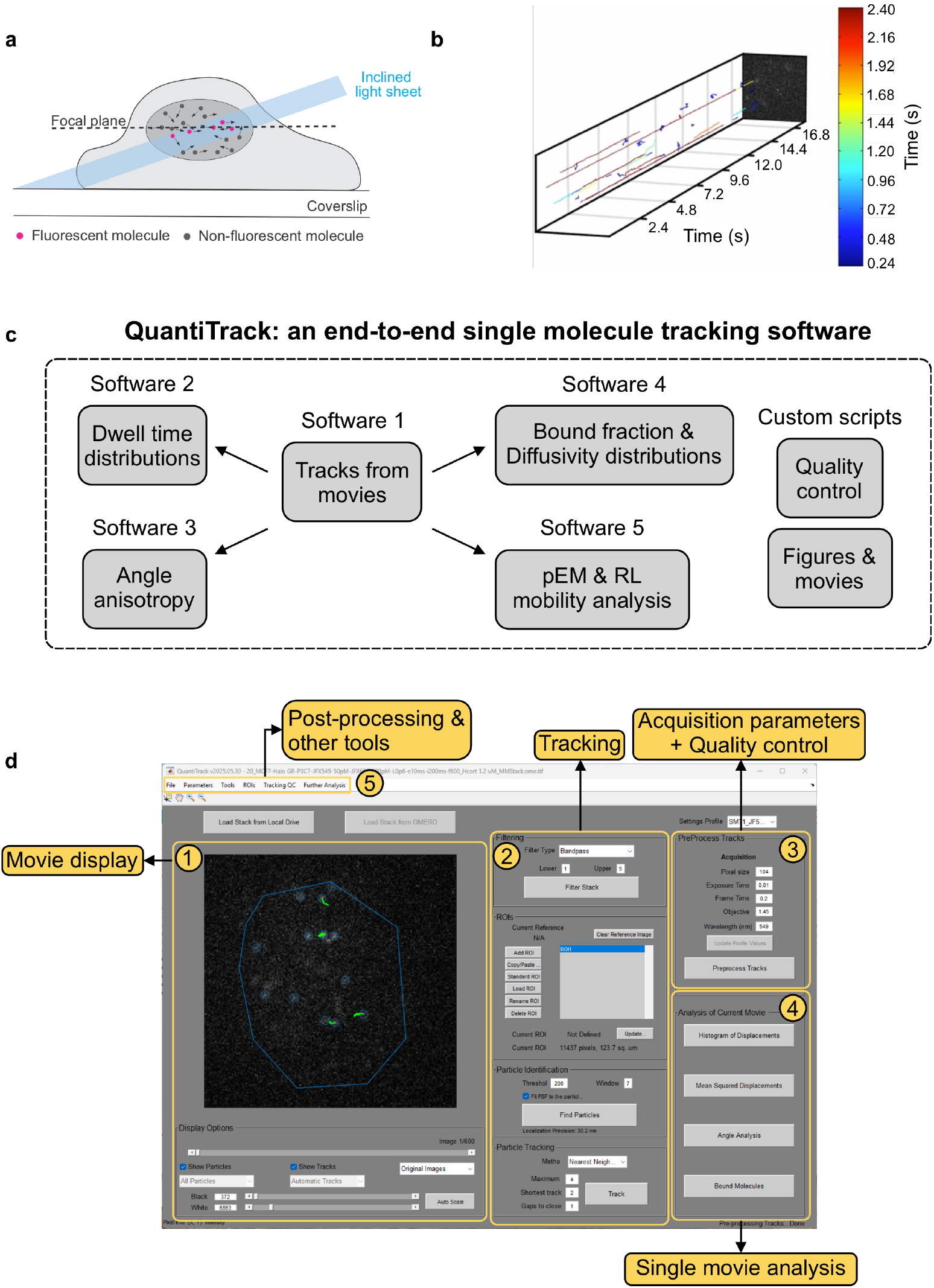
QuantiTrack – a comprehensive GUI based single molecule tracking analysis software. **(a)** Schematic of a single molecule tracking (SMT) experiment. An inclined light sheet is typically used to illuminate a thin section of the cell to minimize background fluorescence. Panel reused with permission from reference^7^ (public domain). **(b)** A (2+1) dimensional snapshot of the output of an SMT experiment. The lines are the detected single molecule trajectories, color-coded according to their track lengths. **(c)** A typical SMT analysis workflow involves the use of multiple software packages to obtain tracks, evaluate detection and tracking quality, perform different analyses, and generate movies and publication-quality figures. QuantiTrack provides an end-to-end solution by unifying all these features into a single graphical user interface (GUI)-based software. **(d)** The GUI for QuantiTrack can be divided into five sections: (1) <u>Movie display panel:</u> Display and controls for viewing individual movie frames and detected tracks. (2) <u>Tracking panel:</u> To define the region of interest (ROI), select detection and tracking parameters (3) <u>Acquisition parameters and quality control panel:</u> To set the acquisition parameters and perform quality control for the loaded movie. (4) <u>Single movie analysis panel:</u> This panel contains several options of quick analyses for the loaded movie. (5) <u>Post-processing & other tools panel:</u> Additional analyses and tools are available here.

The QuantiTrack GUI can be divided into 5 panels **(Figure 1d)**:

1. <u>Movie display panel:</u> Here, a user can load in their single molecule movie, adjust visualization parameters and toggle through the entire movie.
2. <u>Tracking panel:</u> The user can select various parameters related to filtering the movie, defining the region of interest (ROI) within the field of view where the particles should be detected, defining particle detection parameters, and methods of particle tracking.
3. <u>Acquisition parameters and quality control panel:</u> Through this panel, the user sets the data acquisition parameters and accesses the QC window through the ‘Preprocess Tracks’ button (see User Guide and **Figure 2** for more details).
4. <u>Single movie analysis panel:</u> This offers four types of analyses for the currently loaded movie.
5. <u>Post-processing & other tools panel:</u> These panels allow the user to perform the analyses shown in **Figure 1c** over all the movies in their datasets.

**Figure 2.**
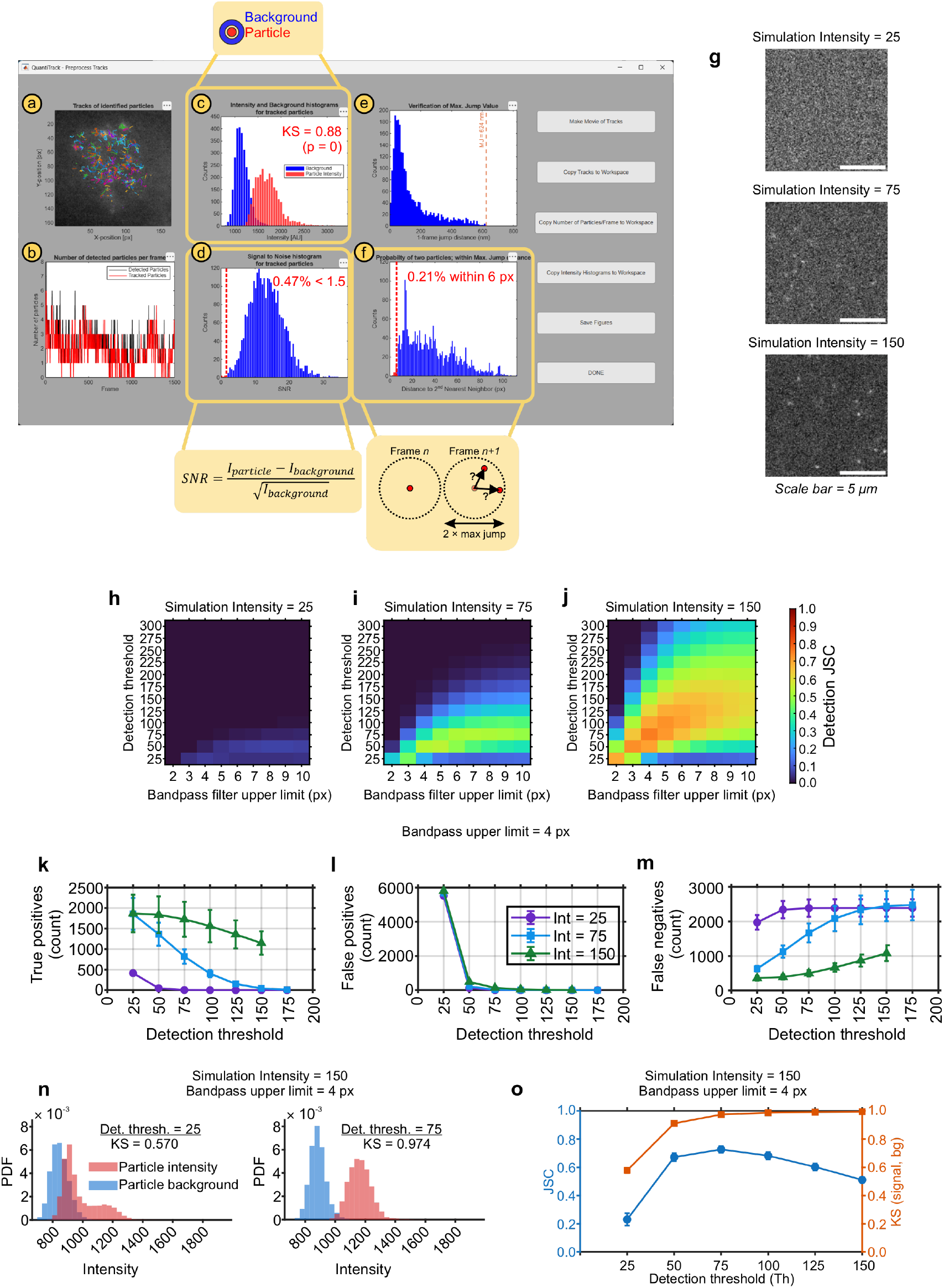
Detection fidelity as a function of signal intensity. **(a–f)** The ‘Preprocess Tracks’ window contains the following qualitative and quantitative measures to assess data quality: **(a)** Tracks overlaid on a temporal projection of the SMT movie. **(b)** Number of detected particles (black) and tracked particles (red) over time, used to assess the fraction of detections incorporated into trajectories based on the chosen tracking parameters. **(c)** Histograms of signal (red) and local background (blue) intensities. (Callout) Particle signal intensity is calculated as the summed intensity of the Airy disk, while the local background is estimated from a surrounding donut region. The text inset is the Kolmogorov-Smirnov (KS) statistic which quantifies the distance between the signal and background intensity distributions. **(d)** Histogram of signal-to-noise ratios (SNR) for all detected particles. Bins with SNR < 1.5 are shown in red to the left of the dashed red line. The inset reports the percentage of particles with SNR < 1.5. (Callout) SNR calculation formula. **(e)** Histogram of single frame displacements with the max jump indicated with the dashed line. **(f)** Histogram of distances between a tracked particle in frame *n* and its second nearest neighbor in frame *n*+*1*. (Callout) Tracking errors can arise if multiple particles fall within the maximum jump radius (MJ). Distances ≤ MJ are shown in red to the left of the dashed red line. The inset reports the corresponding percentage. **(g)** Representative stills from simulations of Brownian motion with D = 1.0 μm^2^/s and signal intensity set to (top) 25, (center) 75, and (bottom) 150. Scale bar = 5 μm. **(h–j)** Heatmaps showing the measured Jaccard similarity coefficient (JSC) for simulation intensities of (h) 25, (i) 75, (j) 150 as a function of bandpass filter upper limit and detection threshold. **(k–m)** The number of (k) true positives (TP), (l) false positives (FP), and (m) false negatives (FN) detected at different detection thresholds. Error bars = standard deviation across 10 movies. The bandpass filter upper limit was set to 4 pixels. **(n)** Signal (red) and local background (blue) intensity histograms for simulation intensity = 150 and detection threshold = 25 (left) and 75 (right). Inset text is the Kolmogorov-Smirnov (KS) statistic between the signal and background distributions. **(o)** JSC (left axis) and KS statistic (right axis) as a function of the detection threshold (simulation intensity = 150, bandpass upper limit = 4 px).

Before analyzing SMT movies, we recommend that each movie be visually inspected for cell-scale motion. Movies exhibiting visible linear drift or rotation should be discarded before tracking because these movements will obscure any molecular motion **(Supplementary Figure 1b)**. Once the movies have been inspected, a typical QuantiTrack workflow involves loading the movie into the GUI; applying top-hat, Wiener, and bandpass filters (see Methods); defining the ROI; iteratively detecting and tracking particles while using the quality control metrics to select the optimal detection and tracking parameters **(Supplementary Figure 1a)**. The next section describes QC metrics available within QuantiTrack, and how to use them to determine optimal particle detection and tracking parameters.

### Using the quality control metrics within QuantiTrack to optimize particle detection

**Figure 2a–f** shows a representative screenshot of the pop-up window that appears after clicking the ‘Preprocess Tracks’ button in the main QuantiTrack GUI (**Figure 1d, panel 3**). This window allows the user to inspect the quality of particle detection and tracking and change any parameters if needed. The first panel **(Figure 2a)** contains an overlay of all the tracks over a time-projection of the SMT movie providing a qualitative overview of the track distributions. **Figure 2b** shows the number of detected particles (black) and the number of tracked particles (red), allowing the user to gauge the fraction of detected particles that are linked to tracks with the current settings. Some of the detected particles may not be tracked – for example, particles that produce tracks shorter than the ‘shortest track’ parameter **(Figure 1d, panel 2)** will not be recorded as tracks.

Two quantitative measures to evaluate particle detection fidelity are presented in **Figure 2c–d**. For every detected particle, the particle intensity is calculated from the total intensity within the Airy disk surrounding the particle coordinates, and the local background is calculated from a donut shaped ROI around this Airy disk (**Figure 2c callout**). **Figure 2c** shows an example of the signal (particle intensity) and noise (background) histograms when the particle detection is robust (see *“Using the quality control metrics within QuantiTrack to optimize particle detection”*). The inset text shows the Kolmogorov-Smirnov (KS) statistic, which is a measure of the separation between the signal and background intensity distributions. QuantiTrack also measures the SNR for all the detected particles **(Figure 2d, callout)** and calculates the fraction of particles with SNR ≤ 1.5 (**Figure 2d**; see Methods), which can be used to measure the fraction of low SNR detections.

Particle detection is susceptible to two types of errors: (i) A false positive (FP) occurs when a detection is registered where no *bona fide* signal exists, and (ii) a false negative (FN) occurs when a particle is not localized despite the presence of signal **(Supplementary Figure 2a)**. In this section, we highlight how the choice of the detection threshold parameter (**Figure 1d, panel 2)** influences both FP and FN errors. We employ simulations to demonstrate how signal intensity (which is typically controlled by the excitation laser power) affects detection fidelity. For practical purposes, we show how the KS statistic can be used to determine the optimal detection threshold parameter.

We used sptPALMsim^29^ (an optical-dynamic simulation package) to simulate Brownian motion with a diffusion coefficient of 1 μm^2^/s and varying signal intensity. Low, intermediate, and high intensity simulations correspond to simulation intensities of 25, 75, and 150 respectively **(Figure 2g and Movie 2)**. Any detection that falls within 4 pixels (416 nm) of a ground truth (GT) localization is classified as a true positive (TP). FP and FN are defined as above **(Supplementary Figure 2a)**. The Jaccard similarity coefficient (JSC), computed as the ratio of TP to the sum of TP, FP, and FN is used to measure detection fidelity^30^.

We next varied both the bandpass filter upper limit (h_pass_) along with the detection threshold and calculated the JSC for all three intensity simulations. The detection threshold parameter determines which local maxima in each filtered image are retained as candidates for PSF fitting. For each combination of detection threshold and h_pass_, the JSC increased with the simulation intensity **(Figure 2h–j)**. The highest detection fidelity is observed for the high intensity simulation with detection threshold = 75 and h_pass_ = 4 px **(Figure 2h–j)**. In our experience h_pass_ = 4 or 5 pixels works well for most experimental datasets.

Since the detection threshold has a larger effect on the JSC than h_pass_, we next examined how this parameter regulates TP, FP, and FN errors with h_pass_ = 4 pixels. For all three datasets, the number of TP and FP decrease monotonically as a function of the detection threshold **(Figure 2k–l)**. Increasing the detection threshold initially reduces FP **(Figure 2l)** but beyond the optimal value also removes lower intensity TP detections **(Figure 2k)**. On the other hand, increasing the detection threshold makes particle detection more stringent thereby reducing the number of FN **(Figure 2m)**.

The optimal JSC, while a useful metric when comparing to the GT, is not practical for an experimental dataset where the GT is inaccessible. Increasing the detection threshold serves to increase the separation between the signal and local background intensity histograms **(Figure 2n, Supplementary Figure 2b–d)**. The optimal detection threshold thus marks the point at which no substantial further separation between the two histograms can be achieved by increasing the detection threshold. The distance between the two histograms can be quantified using the KS statistic (see Methods). While the JSC peaks at the optimal detection threshold, the KS statistic first increases and then reaches a plateau **(Figure 2o, Supplementary Figure 2e–g)**. This elbow or inflection point is precisely the point after which no significant separation is achieved by further increasing the detection threshold. Thus, by measuring the KS statistic for a few representative movies over a range of detection thresholds, users can determine the optimal detection threshold as follows:

1. Sweep the detection threshold in steps of 25 or 50 over a broad range.
2. For each detection threshold, record the KS statistic from the Preprocessing window **(Figure 2c)**.
3. Identify the inflection point in the KS statistic vs detection threshold plot and use this as the detection threshold.

The detection threshold determined from 2–3 representative movies can be applied for other movies with comparable expression, labeling density, and acquisition parameters.

### The effect of motion blur on detection fidelity

During the finite exposure time of the camera, a rapidly diffusing particle’s fluorescence gets blurred over the space it travels. This motion blur reduces the localization precision of detected molecules. Shorter exposure times along with a concomitant increase in the laser power can mitigate this to some extent, but every microscope has a limit to the acquisition speed. To quantify the effect of motion blur on detection accuracy, we simulated normal Brownian motion (simulation intensity = 150) across diffusion coefficients spanning 1 to 20 μm^2^/s, with **(Movie 3)** and without **(Movie 4)** motion blur. The degree of motion blur is determined by the combination of diffusion coefficient and exposure time **(Figure 3a)**. The camera exposure time was set to 10 ms with 12 ms intervals (see Methods), matching our microscope’s fastest acquisition speed and providing a tradeoff between collecting sufficient photons for reliable particle detection while minimizing motion blur. At lower diffusion coefficients, both the motion blur (MB) and no motion blur (NoMB) images show punctate spots corresponding to single molecules. At higher diffusion coefficients, however, the MB images do not retain these sharp punctate spots **(Figure 3a, Movies 3–4)**, illustrating the blurring of fast-moving molecules.

**Figure 3.**
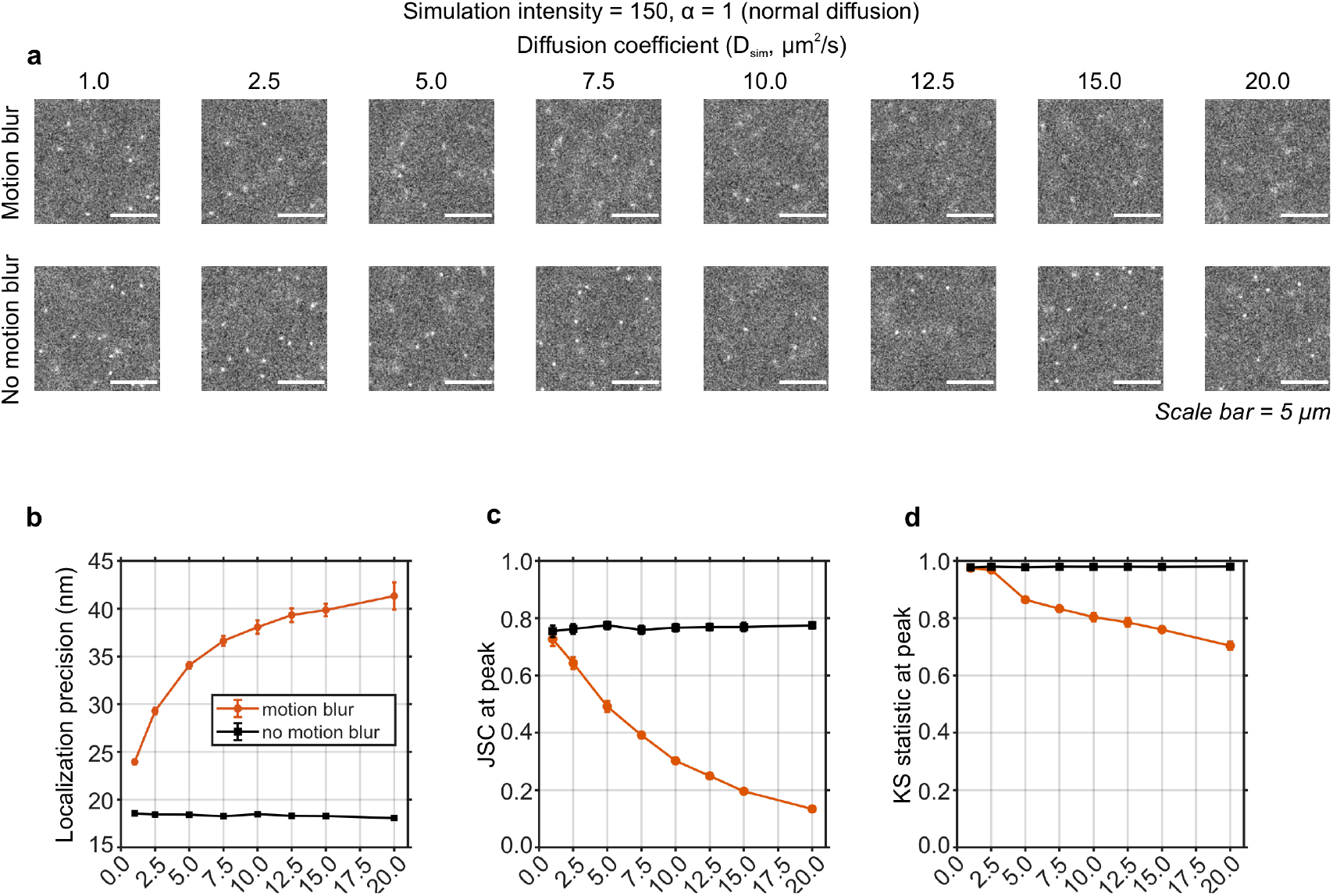
Effect of diffusion coefficient on detection fidelity. **(a)** Representative stills from movies simulated with normal diffusion (α = 1) and indicated diffusion coefficients. (Top row) Simulations with motion blur. (Bottom row) Simulations without motion blur. **(b)** Measured localization precision (in nm) in the presence (orange circles) and absence (black squares) of motion blur. **(c–d)** (c) Jaccard similarity coefficient (JSC) and (d) Kolmogorov-Smirnov (KS) statistic at optimal detection thresholds for simulations with (orange circles) and without (black squares) motion blur.

We next performed the detection threshold sweep as described in “*Using the quality control metrics within QuantiTrack to optimize particle detection”* and identified the optimal detection threshold for each of these datasets. The localization precision, JSC, and KS statistic were then computed for the MB and NoMB datasets. In the absence of motion blur, the diffusion coefficient does not affect the localization precision. On the other hand, motion blur degrades the localization precision from 24 nm to 41 nm **(Figure 3b)**, JSC from 0.72 to 0.13 **(Figure 3c)**, and the value of the KS statistic at the respective inflection points from 0.97 to 0.7 **(Figure 3d)**. Thus, motion blur constrains the range of diffusion coefficients that can be reliably tracked by reducing the detection efficiency. The JSC reduces to 50% of its maximum value for D = 7.5 μm^2^/s. Reliable detection of faster diffusing molecules therefore requires shorter exposure times. Because higher diffusion coefficients also result in larger displacements, and potential tracking errors, we next discuss how diffusion coefficient and labeling density affect tracking fidelity.

### Optimizing tracking fidelity

After optimizing the detection parameters, the next step is to link the individual localizations into trajectories. QuantiTrack offers two tracking algorithms. The first is based on a global nearest-neighbor (NN) assignment subject to a maximum jump (MJ) constraint while the second is u-track’s Linear Assignment Problem (LAP) tracker^26^. While we benchmark the nearest neighbor algorithm here, the User Guide contains further information about the LAP tracker.

Three parameters can be tuned in the nearest-neighbor tracker **(Figure 1d, panel 2)**: (i) the maximum jump parameter ([MJ], pixels), (ii) the shortest track ([ST], frames), and (iii) gaps to close (frames). MJ specifies the largest allowed displacements, ST defines the minimum number of localizations required to register a track, and the gap parameter allows for short gaps caused due to fluorophore blinking. Here, we focus on the role of the MJ parameter in tracking accuracy.

To determine how MJ should be adjusted for different diffusion coefficients, we analyzed the normal diffusion simulations described in the previous section with D = 1–10 μm^2^/s **(Figure 4a)**. Since motion blur precludes accurate detection of particles diffusing faster than 10 μm^2^/s for our choice of exposure time and imaging interval **(Figure 3c)**, we limit our analysis to this range. We varied MJ from 4 pixels (416 nm) to 16 pixels (1664 nm) and measured the tracking accuracy.

**Figure 4.**
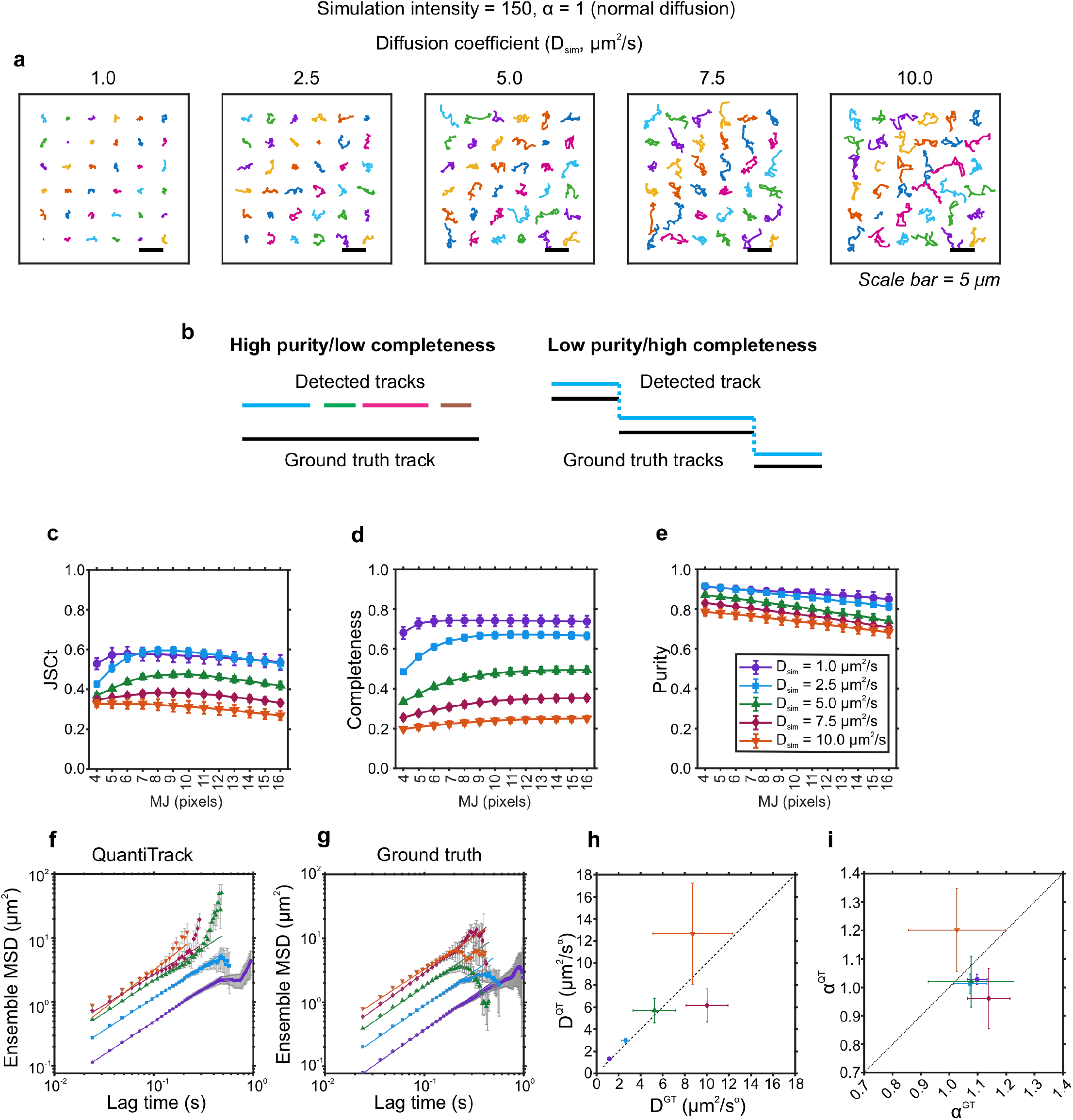
Tracking fidelity as a function of diffusion coefficient. **(a)** Representative trajectories for simulations of normal diffusion with the indicated diffusion coefficient. **(b)** Illustrative examples of (left) high purity/low completeness and (right) low purity/high completeness trajectories. **(c–e)** Measured (c) track-level Jaccard similarity coefficient (JSCt), (d) completeness, and (e) purity as a function of the maximum jump (MJ) parameter for the indicated diffusion coefficients. **(f–g)** Ensemble mean squared displacement (MSD) for (f) tracks identified using QuantiTrack (QT) and (g) ground truth (GT) tracks. Power-law fits (MSD ∼ 4D_α_t_α_) are shown in solid lines. Error bars = s.e.m. **(h–i)** QuantiTrack-estimated vs ground truth (h) diffusion coefficient (D_α_), and (i) MSD power-law exponent (α). Error bars = 95% confidence intervals from the power-law fits.

Particles undergoing Brownian motion exhibit jump distances described by the Rayleigh distribution. To capture the complete range of displacements, MJ should be sufficiently large such that the probability of observing jumps larger than MJ is negligible. To illustrate this, we calculated the jump distance distributions for all the simulated diffusion coefficients at MJ ranging from 4 to 16 pixels and contrasted these with the GT jump distance distributions. For D = 1 μm^2^/s, MJ = 4 px (416 nm) is sufficient to capture most of the displacements **(Supplementary Figure 3a)**. As the diffusion coefficient increases, larger MJ values are required to capture the entire range of displacements **(Supplementary Figure 3b–e)**. The Preprocessing window shows the jump distance histograms for the selected tracking parameters and can be used to optimize the MJ parameter **(Figure 2e)**. Abrupt drops in the jump distance histograms indicate that MJ is too short to capture some of the faster diffusing species **(Supplementary Figure 3b–e)**.

To measure the tracking accuracy, we compute three metrics: (i) the track-level JSC (JSCt), which is the track-level analog of the detection JSC (see Methods), (ii) completeness, and (iii) purity (see Methods). Completeness is defined as the fraction of a GT track’s localizations that map to a single QuantiTrack track. Purity measures the fraction of QuantiTrack track localizations that map to a single GT track. For example, if QuantiTrack detects several short tracks that each completely map onto a GT track, that would be an example of high purity (each of the QuantiTrack tracks completely map to a single GT track) but low completeness (only a small fraction of the GT track localizations map onto a single QuantiTrack track). Conversely, if one QuantiTrack track spans multiple GT tracks, that would be an example of low purity and high completeness **(Figure 4b)**.

For D = 1–7.5 μm^2^/s, JSCt increased with MJ before reaching a plateau, while JSCt for D = 10 μm^2^/s initially remained constant and then decreased slightly with MJ **(Figure 4c)**. As seen from the completeness and purity metrics, the higher diffusion coefficient trajectories were more likely to be fragmented (low completeness) than jump between distinct particles (high purity, **Figure 4d– e**). The higher MJ allowed for more of the larger jumps to be captured **(Supplementary Figure 3b–d)**. At D = 10 μm^2^/s, even at MJ = 16 px (1664 nm), the probability of observing jumps longer than MJ was non-zero **(Supplementary Figure 3e)**. Low MJ values thus truncate trajectories containing displacements larger than MJ, which reduces the completeness.

We computed the ensemble MSDs and fit them with a power-law model for the QuantiTrack trajectories with MJ = 16 px and GT trajectories **(Figure 4f–g)**. The analysis showed that D ≤ 5 μm^2^/s could be accurately recovered, indicating the upper limit for the diffusion coefficients that can be measured under the simulated imaging interval, exposure time, and labeling density **(Figure 4h)**. The localization precision sets the lower limit of diffusion coefficients that can be faithfully measured. If σ is the localization precision, 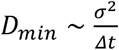. Thus, for σ ∼ 30 nm and Δt = 12 ms, D_min_ ∼ 0.07 μm^2^/s is the smallest resolvable diffusion coefficient under these acquisition parameters. The scaling exponent α, was recovered to within 10% of its input value for all D except D = 10 μm^2^/s **(Figure 4i)**.

High MJ values are only useful if the larger MJ does not incorrectly link particles detected in consecutive frames. For a particle in frame *n*, the presence of two or more particles within MJ in frame *n*+*1* could create erroneous links and increase the chance of mistracking. This can be quantified by measuring the distance between every tracked particle and its second nearest neighbor in the subsequent frame (see Methods). From the histogram of all such distances, QuantiTrack calculates the fraction of second nearest neighbors which fall within MJ **(Figure 2f)**. When tracking fast diffusing particles, we suggest ensuring that both the probability of observing jump distances longer than MJ is negligible **(Figure 2e and Supplementary Figure 3)** as is the fraction of second nearest neighbors within MJ **(Figure 2f)**. A high fraction of second nearest neighbors within MJ indicates that the labeling density is too high, and this should primarily be mitigated experimentally, by reducing the labeling density (see Discussion).

To provide a benchmark of the effect of labeling density on tracking accuracy, we varied the labeling density from 10 to 90 particles/frame for D = 1 and 5 μm^2^/s **(Supplementary Figure 4)**. For D = 1 μm^2^/s, the tracking accuracy did not degrade substantially up to a density of 30 particles/frame **(Supplementary Figure 4a–d)**. However, for D = 5 μm^2^/s, even an increase from 10 to 20 particles per frame resulted in a measurable reduction in JSCt, largely driven by a loss of completeness **(Supplementary Figure 4e–h)**.

Thus, the labeling density as well as the MJ parameter need to be optimized based on the expected motion of the protein of interest. For each new protein and cell line combination, these parameters should be optimized on a representative set of movies and subsequently kept fixed across movies acquired for the same protein, cell line, dye, treatment, and acquisition parameters.

In the next two sections, we use simulations to demonstrate the analyses available in QuantiTrack.

### Deciphering mobility states in multicomponent systems

Molecules navigating the heterogenous intracellular environment can exhibit diverse modes of motion. QuantiTrack includes three analyses for identifying the distinct mobility states within a set of trajectories **(Supplementary Table 1)**. To illustrate these approaches, we simulated normal diffusion (α = 1) with varying proportions of slow and fast diffusing molecules (D_slow_ = 0.1 μm^2^/s and D_fast_ = 2.5 μm^2^/s, **Figure 5a)**.

**Figure 5.**
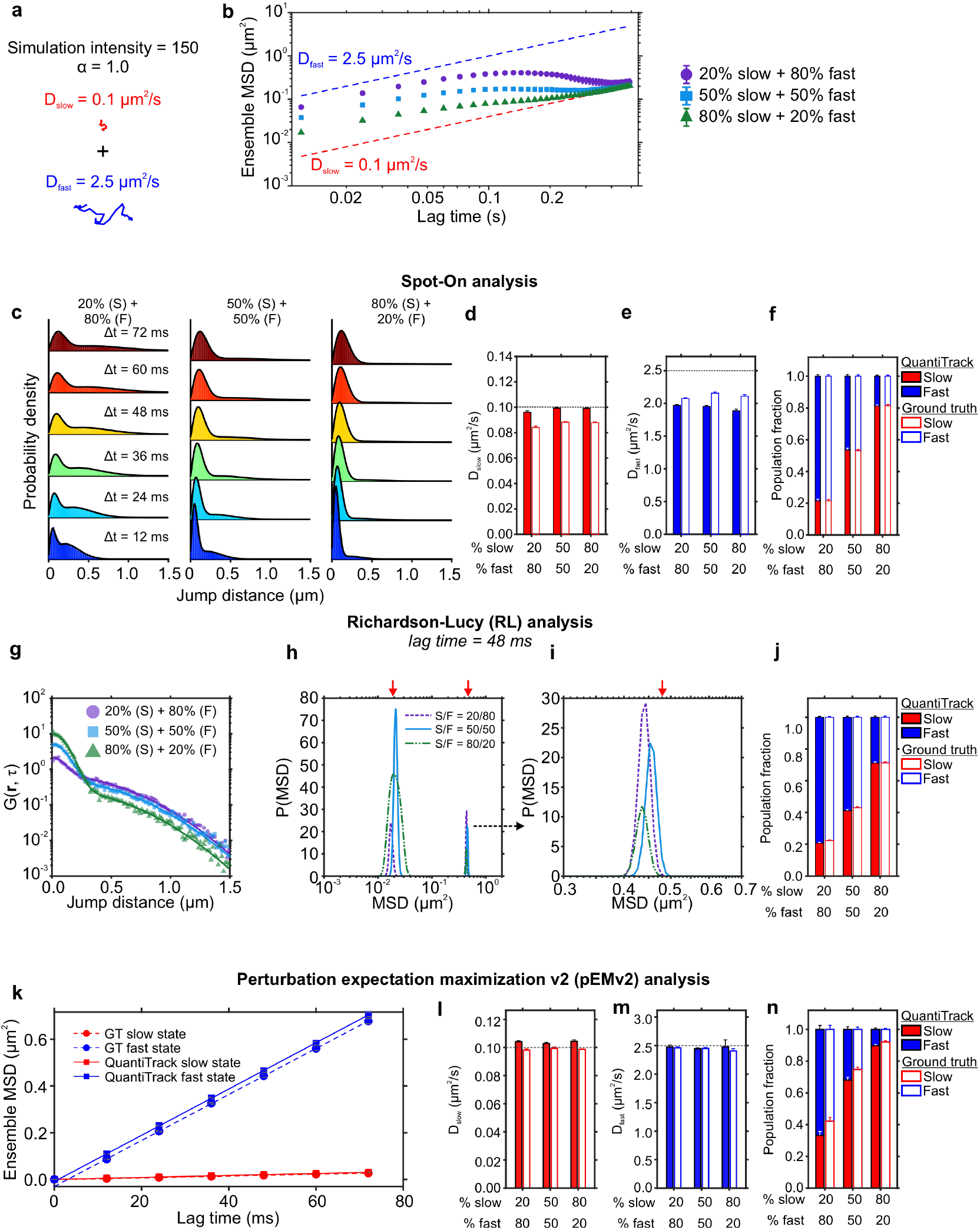
Inferring mobility states from a mixed population. **(a)** Two states with varying proportions of slow and fast diffusion are simulated with D_slow_ = 0.1 μm^2^/s and D_fast_ = 2.5 μm^2^/s, each with α = 1.0. **(b)** Ensemble mean squared displacement (MSD) for indicated mixtures of slow (S) and fast (F) states. Dashed lines indicate the MSD for each of the individual states. Error bars = s.e.m. **(c)** Jump distance histograms and model fits from a two-state model using Spot-On are shown for the indicated mixtures of slow and fast states. **(d–f)** Inferred (d) D_slow_, (e) D_fast_, and population fractions from the QuantiTrack and ground truth datasets with indicated proportions. Error bars = standard deviations of 100 bootstrapped runs. **(g)** van Hove correlation at lag time = 48 ms and model fits using the Richardson-Lucy (RL) deconvolution for the three slow/fast mixtures. **(h)** MSD distribution inferred from the RL analysis. Red arrows indicate the true MSD of the slow and fast states at lag time = 48 ms. **(i)** Zoom-in view of the high MSD peaks from h. **(j)** Population fractions of the slow and fast states for the QuantiTrack and ground truth trajectories. **(k)** Ensemble MSD for the two states obtained from perturbation expectation maximization v2 (pEMv2) analysis for QuantiTrack (solid) and ground truth (dashed) datasets. Error bars = s.e.m. **(l–m)** Inferred (l) D_slow_ and (m) D_fast_ from the pEMv2 analysis. Error bars = 95% confidence intervals. **(n)** Population fractions of the slow and fast states for the three mixtures. Error bars = bootstrapped standard deviation. N_tracks_ for the QuantiTrack datasets = 16,658 (20% slow + 80% fast), 12,702 (50% slow + 50% fast), and 8,699 (80% slow + 20% fast). N_tracks_ for the ground truth datasets = 19,898 (20% slow + 80% fast), 14,567 (50% slow + 50% fast), and 9,536 (80% slow + 20% fast).

The ensemble MSD for the three mixtures (slow | fast = 20% | 80%, 50% | 50%, 80% | 20%) illustrates a limitation of conventional MSD analysis. While this approach can estimate mobility parameters in a single state system **(Figure 4f–i)**, it cannot resolve individual mobility states within a heterogeneous population of trajectories **(Figure 5b)**.

The first approach, Spot-On^5^, jointly fits the jump distance histograms across multiple lag times using a user-specified two- or three-state model. The resulting fit parameters provide estimates of the diffusion coefficient and population fraction of each state. We applied a two-state model to each mixture across six lag times **(Figure 5c–f)** and compared the resulting parameter estimates with those obtained from their respective GT trajectories (see Methods, **Supplementary Figure 5a–c**). The estimated diffusion coefficients and population fractions were consistent between the QuantiTrack and GT trajectories for all three slow/fast mixtures **(Figure 5d–f)**, However, D_fast_ was underestimated in both datasets, suggesting a potential limitation of Spot-On in estimating the diffusion coefficients of fast states under these conditions **(Figure 5e)**.

Spot-On is useful for estimating diffusion coefficients and population fractions when the number of mobility states is known. However, this is often precisely the information that needs to be inferred from the data. QuantiTrack includes a second approach that estimates the distribution of MSDs (and thus, the distribution of diffusion coefficients) without requiring *a priori* knowledge of the number of states. This method is based on an iterative scheme inspired by the Richardson-Lucy algorithm for image deconvolution, and has been used previously to analyze single molecule trajectories^9,31^. This method is referred to as ‘RL analysis’ in QuantiTrack (see User Guide, Methods). We used this method to estimate the MSD distribution from the van Hove correlation (vHc) or jump distance distribution as has been done previously^7,9,25^ **(Figure 5g, Supplementary Figure 5d– f)**. We applied RL analysis to the three mixtures and found distinctly bimodal MSD distributions with the two peaks corresponding to those expected for D_slow_ and D_fast_ **(Figure 5h–i)**. Applying the same analysis to the corresponding GT trajectories produced MSD distributions that closely agreed with those obtained from the QuantiTrack trajectories **(Supplementary Figure 5g–i)**.

Spot-On aggregates jump distances across all trajectories and provides population level parameter estimates but does not classify individual trajectories to the different mobility states. In contrast, following RL analysis, the minima between the peaks in the inferred MSD distributions can be used to classify tracks into distinct mobility groups as demonstrated previously^7,25^ (**Supplementary Figure 5g–l**; see Methods). The resulting population fractions were consistent between both the QuantiTrack and GT datasets (see Methods, **Figure 5j**).

While Spot-On and RL can extract information from the overall jump distributions, they do not yield information about particle mobility states at the level of individual trajectories. QuantiTrack includes an algorithm called perturbation expectation maximization version 2^32^ (hereon referred to as pEMv2) which classifies the motion of trajectories, while accounting for localization errors and motion blur artifacts inherent to SMT **(Supplementary Figure 6a)**. pEMv2 evaluates models containing several mobility states (defined by the user; typically, between 1 and 15) and uses a Bayesian Information Criterion (BIC) to determine the optimal number of states that best describes the collection of trajectories. Given the computational cost associated with inverting large co-variance matrices, long trajectories are split into shorter sub-tracks. Each sub-track is then assigned to the state for which it has the highest posterior probability **(Supplementary Figure 6a)**, thus recovering mobility information at the level of individual trajectories. We empirically determined that seven frame long sub-tracks balance computational cost and classification accuracy^24^ for a large family of TFs for the imaging conditions we have used. A more principled discussion on the choice of sub-track length can be found in the original publication^32^.

When pEMv2 was applied to the GT and QuantiTrack trajectories, only two statistically significant mobility states (with the appropriate diffusion coefficients) were found for all three mixtures **(Supplementary Figure 6b–d)**. Since each sub-track is assigned to a state, ensemble MSDs can be calculated for each state and the diffusion coefficient inferred by fitting the MSDs to a diffusion model **(Figure 5k and Supplementary Figure 6c)**. The estimated diffusion coefficients were consistent between the QuantiTrack and GT datasets and matched the simulation input D_slow_ and D_fast_ **(Figure 5l–m)**. While the population fractions were also consistent between the QuantiTrack and GT datasets, the slow state population fraction was overestimated for all three mixtures **(Figure 5n)**. This discrepancy may arise from differences in how pEMv2 calculates the population fractions compared to those calculated from Spot-On and RL analysis (see Discussion and Methods).

QuantiTrack thus provides three complementary approaches to identify distinct mobility states in complex biological data, each with their own strengths and limitations (see Discussion).

### Turning angle anisotropy as a measure of spatial exploration

While the MSD measures how the spatial extent of a trajectory changes with time, the turning angle distribution can be used to examine the directional persistence of the trajectories^16,17,33,34^. To illustrate this, we simulated three types of fractional Brownian motion (fBM), each with D_α_= 1 μm^2^/s_α_ and scaling exponent α = 0.7 (sub-diffusion), 1.0 (normal diffusion), and 1.3 (super-diffusion) **(Figure 6a–d)**.

**Figure 6.**
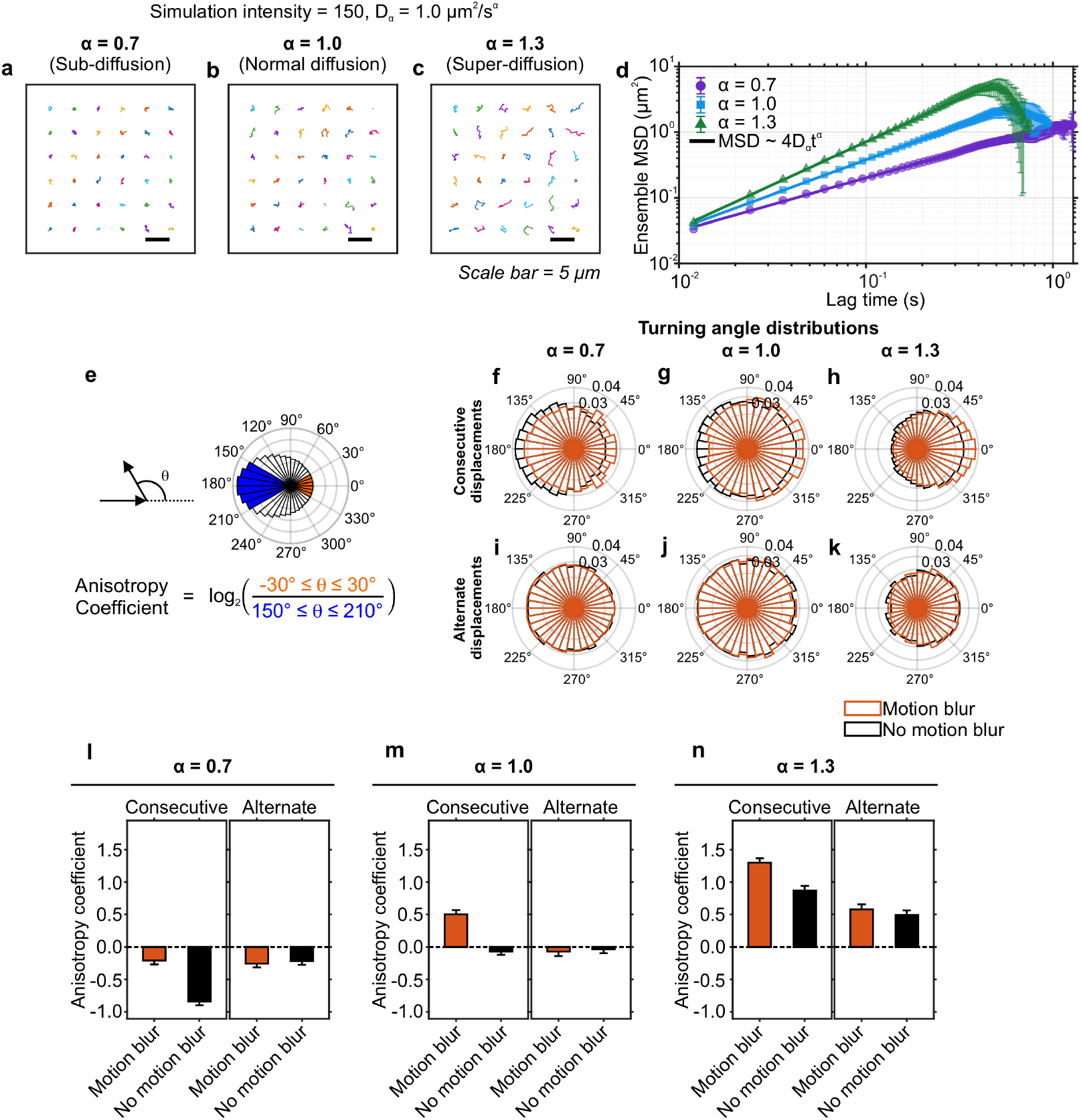
Turning angle anisotropy and the effect of motion blur. **(a–c)** Representative trajectories of simulated fractional Brownian motion with α = (a) 0.7, (b) 1.0, (c) 1.3. D_α_ = 1.0 μm^2^/s_α_ for all three. **(d)** Ensemble mean squared displacement (MSD) and power-law fits for the three datasets. Error bars = s.e.m. **(e)** (Top left) The turning angle θ is the angle between two displacement vectors. (Top right) Polar histogram of turning angles with forward displacements indicated in orange and backward displacements in blue. (Bottom) Anisotropy coefficient (AC) is calculated as the log_2_ ratio of the probabilities of forward (orange) to backward (blue) jumps. **(f–k)** Polar histogram of turning angles for indicated α. (f–h) Turning angles between consecutive displacements (e.g. 1®2 and 2®3). (i–k) Turning angles between alternate displacements (e.g. 1®2 and 3®4). Orange = simulations with motion blur, black = simulations without motion blur. **(l–n)** Anisotropy coefficient calculated for consecutive and alternate displacements for indicated α. Error bars = bootstrapped standard deviation.

The angle between two displacement vectors is called the turning angle (θ). Forward jumps are defined as −30° ≤ *θ* ≤ 30° while backward jumps are defined as 150° ≤ *θ* ≤ 210° **(Figure 6e)**. The turning angle anisotropy coefficient (AC) is defined as the log_2_ ratio of the probabilities of forward and backward jumps^16^ **(Figure 6e)**. Variations of this metric have been previously described, for example in references^17,34^. Negative AC values indicate more confined tracks, pure Brownian motion corresponds to AC = 0, while directional motion and super-diffusion correspond to AC > 0.

While consecutive displacements may appear correlated due to motion blur, alternate displacements (e.g. displacements 1®2 and 3®4) are immune to the motion-blur induced covariance **(Supplementary Figure 7)**. True anisotropy should persist beyond the covariance introduced between successive displacements by motion blur. We measured the turning angles between alternate displacements and found no significant difference between the MB and NoMB trajectories within any of the three α regimes **(Figure 6i–k)**.

These observations can be quantified using the AC. For consecutive displacements, motion blur increases the forward bias (and therefore shifts AC towards more positive values) for all three α. For alternate displacements, AC is not affected by motion blur **(Figure 6l–n)**. The negative AC for sub-diffusive trajectories indicates an increased probability of backward displacements relative to forward displacements **(Figure 6l)**, while the positive AC for super-diffusive trajectories indicates a bias towards forward displacements **(Figure 6n)**. For normal diffusion, AC ∼ 0 in the absence of motion blur, which implies that the directions of consecutive displacements are independent of each other **(Figure 6m)**. While motion blur produced a positive AC for normal diffusion, this bias vanished when alternate displacements were analyzed **(Figure 6m)**.

While angular anisotropy and turning angle distributions are useful tools for investigating how a protein navigates the cellular microenvironment, these measurements alone do not suffice to describe their motion. Quantitative modeling and simulations should be used to completely interpret the angular anisotropy^16,35^ (see Discussion).

To demonstrate the utility of QuantiTrack, we next applied it to quantitatively study GR dynamics under conditions of hormone treatment and washout.

### Glucocorticoid receptor target genes and GR dynamics respond to hormone treatment and washout

GR is activated upon the binding of its cognate ligands (glucocorticoids [GC]) and serves as a TF that regulates the stress response, inflammation, metabolism, cognitive function, and immune response, among others^36^. GR is primarily cytoplasmic in the absence of hormone, sequestered by a chaperone complex. Upon ligand binding, GR translocates to the nucleus and remains nuclear after hormone washout^20,37^. When conjugated to an agonist, GR dynamically binds glucocorticoid response elements within enhancers as a tetramer^38^ **(Figure 7a)**.

**Figure 7.**
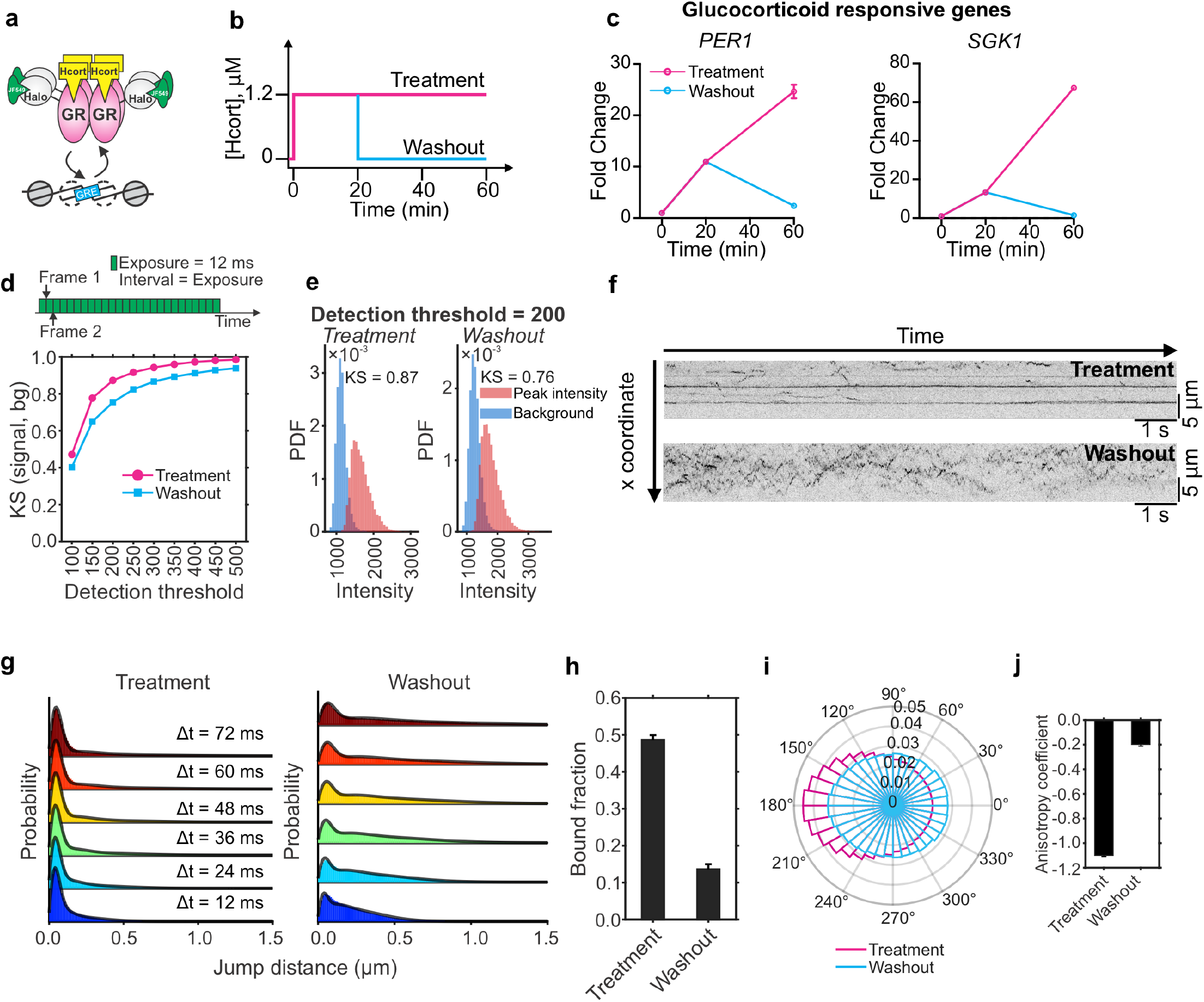
Hormone washout leads to downregulation of GR-responsive genes and significantly reduces GR bound fraction. **(a)** Activated GR dynamically binds to glucocorticoid response elements (GREs) as a tetramer. **(b)** Schematic of hormone treatment and washout protocols. Hcort = hydrocortisone, a natural glucocorticoid. **(c)** qPCR data for two GR responsive genes (*PER1* and *SGK1*) under Hcort treatment and washout at the indicated time points. Error bars = s.e.m. All data are normalized to the vehicle control. **(d)** (Top) Schematic of the imaging protocol for fast SMT. Images are collected every 12 ms with continuous exposure for 1500 frames. (Bottom) Kolmogorov-Smirnov (KS) statistic for the signal and local background intensity distributions as a function of detection threshold. Magenta circles = GR + Hcort treatment and cyan squares = GR + Hcort washout. **(e)** Representative signal (red) and background (blue) intensity distributions for (left) treatment and (right) washout conditions. The KS statistic is reported in the inset text. **(f)** Representative kymograms of GR under Hcort treatment (top) and Hcort washout (bottom). Stripes indicate bound molecules. **(g)** Jump distance distributions across six lag times along with three-state Spot-On model fits (solid lines) for (left) GR + Hcort treatment and (right) washout. **(h)** Bound fraction measured from Spot-On. Error bars = standard deviation across 100 bootstrapped samples. **(i)** Polar histogram of displacement angles for Hcort treatment (magenta) and washout (cyan). **(j)** Anisotropy coefficient for the polar histograms shown in panel i. Error bars = bootstrapped standard deviation across 500 resamples. N_cells_/N_tracks_ = 130/93,836 (GR + Hcort-treatment) and 133/108,696 (GR + Hcort-washout)

In mammals, GCs are secreted in circadian rhythms, with the peak in hormone release coinciding with the active phase of the animal^39^. Underlying this circadian pattern, are short, roughly hourly, ‘ultradian’ pulses of glucocorticoid release^40^. Work from several labs, including ours, has studied the effect of pulsatile hormone release on GR-responsive gene expression^20,23,41^. In response to ultradian GC pulses, many GR-target genes are also transcribed in a pulsatile manner^20,21,23,41,42^. The mechanism behind pulsatile transcription has been linked to differential recruitment of GR to GREs^20^ and enhancer-specific changes in accessibility^23^. FRAP experiments showed that compared to constant hormone exposure, GFP-GR fluorescence recovery at a tandem array of GR binding sites is faster upon hormone washout^20^. However, FRAP represents an ensemble measurement, with diffusing molecules being major contributors to the FRAP curves^43^. No detailed single molecule characterization of GR dynamics under hormone treatment and washout conditions has been performed. Here, we use this well-defined system to show how QuantiTrack enables quantitative, single molecule dissection of GR binding dynamics upon hormone stimulation and washout.

To simulate the effects of hormone pulses, we expose cells to a 20 min pulse of hydrocortisone (Hcort), which is the pharmaceutical name of the natural human GC cortisol. Following this pulse, we either maintain the cells in Hcort-containing medium, which we refer to as the ‘treatment’ condition in this manuscript, or wash out the hormone using conditioned medium (**Figure 7b**, see Methods). The latter protocol is referred to as the ‘washout’ condition hereon.

We tested the effect of hormone withdrawal on two GR target genes (*PER1* and *SGK1*) in MCF-7 cells and found that both genes are robustly activated 20 min after hormone treatment and exhibit even higher activation at 60 min under the treatment protocol. On the other hand, upon hormone washout, both *PER1* and *SGK1* return to their baseline expression at 60 min **(Figure 7c)**. This is in line with previous studies in murine cell lines as well as in animal models^20,21,42^.

Based on the FRAP data^20^, changes in GR target gene expression likely reflect upstream changes in GR dynamics. To study this, we generated a clonal MCF-7 cell line with a HaloTag knocked in at the endogenous *NR3C1* locus (which encodes the GR protein) using CRISPR-Cas9 (**Supplementary Figure 8a–b**, see Methods). We also generated an MCF-7 cell line expressing H2B-Halo from the endogenous H2B promoter (**Supplementary Figure 8c**, see Methods). As expected, Halo-GR exhibits predominantly cytoplasmic localization in the absence of hormone, and is primarily nuclear upon hormone treatment, while H2B-Halo is nuclear and correlates with DAPI staining **(Supplementary Figure 8d–e)**.

We first determined the fraction of bound GR molecules in the nucleus. To capture both bound and diffusing molecules, we collected images of Halo-GR under Hcort treatment and washout conditions every 12 ms under continuous illumination **(Figure 7d)**. We used the KS statistic to determine the optimal detection threshold (Th = 200) as described in *“Using the quality control metrics within QuantiTrack to optimize particle detection”* **(Figure 7d–e and Supplementary Figure 9)**. Representative kymograms show the stark difference between GR dynamics within the nucleus under hormone treatment and washout, with long stripes visible in the treatment condition but only short, highly dynamic trajectories visible in the washout condition **(Figure 7f, Supplementary Figure 9b and d, and Movie 5)**.

We used Spot-On to estimate the bound fraction of GR under Hcort treatment and washout. To benchmark the range of bound fractions that can be measured with experimental data, we first used publicly available data kindly provided by Anders Hansen and colleagues^17^ to measure the bound fraction of histone H2B-Halo, the architectural protein Halo-CTCF, the TF Halo-SOX2, and HaloTag conjugated to a nuclear localization signal (Halo-3xNLS) in mouse embryonic stem cells with images collected every 7.5 ms. Since H2B is primarily incorporated into chromatin and Halo-3xNLS has no specific binding sites, these two proteins represent the upper and lower reference points, respectively, for bound fractions that can be measured using Spot-On under these acquisition conditions.

For consistency with the analysis performed in the original publication^17^, we first fit a two-state model (with one bound state and one diffusive state) jointly to jump distance histograms across four lag times. The estimated bound fraction ranged from 0.90 for H2B to 0.16 for Halo-3xNLS **(Supplementary Figure 10a–e)**. The two-state model fit the H2B, CTCF, and SOX2 jump distance histograms reasonably well **(Supplementary Figure 10a–c)**, whereas some deviations between the data and model fit were apparent for Halo-3xNLS **(Supplementary Figure 10d)**.

Extending the analysis to a three-state model (with one bound and two diffusive states) improved the agreement between the model and the Halo-3xNLS data **(Supplementary Figure 10f–j)**. Although the measured bound fractions were lower with the three-state model, the ordering among these four proteins was preserved **(Supplementary Figure 10j)**. Among these datasets, the smallest measurable bound fraction was ∼0.1 (for Halo-3xNLS) **(Supplementary Figure 10j)**.

We next used Spot-On to estimate the bound fraction of GR under Hcort-treatment and washout. Both two- and three-state models were applied to the GR jump distance distributions under both conditions (see Methods). Since the three-state model showed less deviations from the data, especially for the washout condition **(Figure 7g, Supplementary Figure 10k**; see Discussion), we report the bound fraction from the three-state model **(Figure 7h)**. Both the two- and three-state models showed that hormone washout led to a significant reduction in the bound fraction of GR (**Figure 7h, Supplementary Figure 10l–m**), in agreement with previous FRAP data^20^.

Analysis of the turning angles showed that GR under Hcort treatment exhibits a higher probability of backward jumps reflecting the higher bound fraction, while hormone washout results in similar probabilities of forward and backward jumps, consistent with the higher diffusive fraction observed under the washout condition **(Figure 7i–j)**.

While fast SMT allows us to track both bound and diffusive molecules, photobleaching prevents the observation of binding events lasting on the order of seconds. To overcome this limitation, we next modified our imaging protocol to quantify the long-timescale spatiotemporal dynamics of bound GR molecules.

### Hormone washout results in shorter GR dwell times and reduced occupancy of mobility states correlated with transcriptional activity

To examine the dynamics of bound GR molecules, we modified our imaging protocol as follows: images were collected with 10 ms exposure times (to minimize motion blur) and an interval of 200 ms (to ensure that fast diffusing molecules are not imaged), for a duration of 2 min **(Figure 8a, Movie 6)**. We used the KS statistic to determine the optimal detection threshold (Th = 200) **(Supplementary Figure 11)**. This slow SMT protocol allows us to specifically capture the motion of bound molecules. By separating the excitation pulses by 200 ms, we minimize both the probability of capturing diffusing molecules and increase the likelihood of tracking bound molecules over tens of seconds. The binding events of GR molecules are visible as ‘stops’ that appear as stripes on a kymogram **(Figure 8a)** for both hormone treatment and washout. The dwell time distribution of these ‘stopped’ events provides a measure of the binding times of TFs^44^. To quantify the temporal behavior of such bound molecules, we must account for photobleaching, which would otherwise result in the underestimation of the true dwell time distribution: molecules that remain in the focal volume longer than photobleaching permits will disappear not because they leave the focal volume, but due to photobleaching. We refer the reader to a recent publication for a detailed discussion on the concept of photobleaching^2^. A stable protein such as histone H2B has been used as a probe for photobleaching in the context of nuclear proteins^18,45^. We therefore collected H2B-Halo data, using the same imaging protocol and acquisition parameters, as a control for photobleaching.

**Figure 8.**
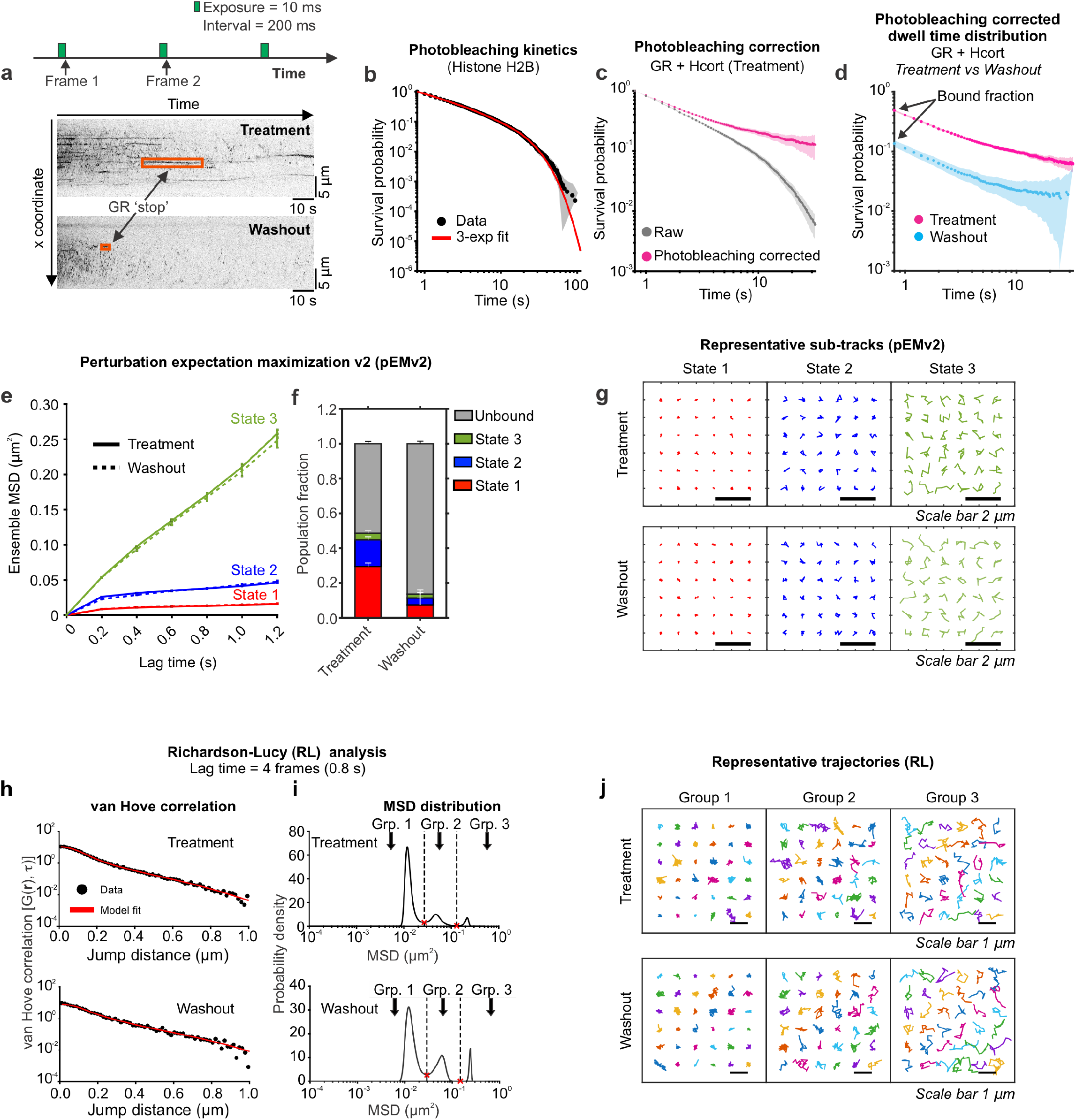
Hormone washout reduces GR dwell times and occupancy of the transcriptionally active mobility state. **(a)** (Top) Imaging protocol for slow SMT. Images are acquired every 200 ms with an exposure time of 10 ms for 2 min (600 frames). (Bottom) Representative kymographs of GR under hydrocortisone (Hcort) treatment (top) and washout (bottom) conditions. Stationary GR molecules appear as stripes (orange boxes). **(b)** Histone H2B survival distribution is used to calculate the photobleaching rate. The raw H2B survival distribution (black) is fit to a triple exponential function (red), and the slowest exponential component defines the photobleaching rate. Shaded error bars indicate 95% confidence intervals (CI). **(c)** Survival distributions for GR under Hcort treatment, before (gray) and after (magenta) correcting for photobleaching. Shaded error bars = 95% CI. **(d)** Photobleaching-corrected survival distributions normalized to the bound fraction for GR + Hcort treatment (magenta) and washout (cyan). Shaded error bars = 95% CI. **(e)** Ensemble mean-squared displacements (MSD) for three mobility states identified using perturbation-expectation maximization (pEMv2). Solid lines: Hcort treatment, dashed lines: Hcort washout. Error bars = s.e.m. N_sub-tracks_ = 18,811 (GR + Hcort treatment), 3,971 (GR + Hcort washout). **(f)** Population fractions of the three pEMv2 states, normalized to the bound fraction. Red: state 1, blue: state, green: state 3, gray: unbound fraction. Error bars = standard deviation of 10,000 bootstrapped samples. **(g)** Representative sub-tracks belonging to each of the three pEMv2 states. (Top) GR + Hcorttreatment and (bottom) washout. Scale bar = 2 μm. **(h)** van Hove correlation for GR + Hcort treatment (top) and washout (bottom). Black circles: data, red: model fit. **(i)** MSD distribution from the RL analysis for GR + Hcort treatment (top) and washout (bottom). The minima in the MSD distribution are used to classify tracks into different mobility groups. **(j)** Representative tracks assigned to the three RL groups for (top) treatment and (bottom) washout. Scale bar = 1 μm. N_cells_/N_tracks_ = 42/33,173 (H2B), 96/58,239 (GR + Hcort treatment), 70/32,183 (GR + Hcort washout).

We first calculated the survival distribution of H2B, which is a key step in accurately measuring the photo-bleaching rate. The survival distribution is the probability of observing a binding event longer than a certain time. Formally, the survival distribution is calculated as *S*(*t*) = 1 − *CDF*(*t*), where the CDF is the cumulative distribution function. We fit the histone H2B survival distribution to a triple exponential function to estimate the photobleaching rate **(Figure 8b)** as has been described previously^7,18,24,46^ (see Methods). The photo-bleaching rate calculated from the H2B data was used to perform the photobleaching correction on the raw GR survival distributions **(Figure 8c)**. Further, the y-intercept of the survival distribution must be normalized to the bound fraction (obtained from the three-state Spot-On model; **Figure 7h**) since the slow SMT protocol only tracks bound molecules. The data showed that hormone washout significantly reduces the binding lifetimes of GR, corroborating our previously published FRAP data^20^ **(Figure 8d)**.

A single TF ‘stop’ that appears as a stripe in a kymogram **(Figure 8a)** is not necessarily a site-specific binding event (see Discussion). Within a single kymogram stripe, the TF molecule engages in different kinds of motion (we refer the reader to a recent review^4^ for a detailed discussion on this). We have previously demonstrated that, on a time scale of 1.2 s, TFs exhibit primarily two distinct mobility states, which are related to the motion of the chromatin fiber^7^. Upon activation with their respective hormones, steroid hormone receptors predominantly bind in the lowest mobility state (which we refer to here as state 1) and binding in this state requires an intact DNA-binding domain as well as domains necessary for the recruitment of transcriptional cofactors^7^.

We applied pEMv2 to GR trajectories under both hormone treatment and washout conditions and found that GR exhibits 3 distinct mobility states: state 1 has the lowest mobility followed by states 2 and 3 **(Figure 8e–g, Supplementary Figure 12)**. After normalizing the pEMv2 fractions to the bound fraction (see Methods), we found that hormone washout resulted in an ∼75% reduction in the fractions of state 1 and state 2 **(Figure 8f)**, suggesting that GR cannot bind effectively in the transcriptionally competent state in the absence of hormone, consistent with previous work using transcriptionally inactive TF mutants^7,24^.

Since pEMv2 operates on sub-tracks, we also applied RL analysis to the slow SMT data to examine the mobility of entire tracks. As we have described previously for GR and other nuclear receptors, RL analysis reveals 3 to 4 distinct mobility groups at a lag time of 0.8 s for both hormone treated and hormone washed out GR molecules^7^ **(Figure 8h–j)**. Note that here we use the term ‘groups’ for the RL groups to avoid confusion with the pEMv2 ‘states’ described above. As before, we used the minima in the MSD distribution to classify the tracks into different mobility groups with increasing exploration areas **(Figure 8i–j)**.

Together, these data show that upon hormone washout, GR bound fraction, dwell times, and the fraction of molecules in the mobility state correlated with transcriptional activation (state 1) drops significantly, which results in the downregulation of GR target genes **(Figures 7 and 8)**.

## DISCUSSION

### QuantiTrack provides an integrated workflow for SMT analysis

Single molecule tracking has become a leading approach for studying protein dynamics and interactions in living cells. Over the past two decades, SMT has been applied to diverse systems – including membrane proteins, molecular motors, and nuclear and nucleolar factors – and has also been used to infer rheological properties of cellular sub-compartments^28^. Despite these advances, broad adoption of this technique by the molecular biology community has been limited, in part, due to the computational expertise required for data analysis.

The basic building blocks of an SMT experiment are a HILO, TIRF, or light sheet microscope equipped with a sensitive camera, along with cells expressing tagged proteins of interest. Due to advances in commercial microscopy platforms, CRISPR-based gene editing, and the widespread availability of organic fluorophores, these components are now increasingly accessible to molecular biology labs. The primary barrier to the adoption of SMT is no longer the instrumentation or reagents but the complexity of data analysis and the challenge of translating SMT results into biological insight.

Many tracking and analysis tools have been developed for specific aspects of the analysis workflow. Switching between tools often requires reformatting data or rewriting code. This underscores the need for an integrated software package that can not only track single molecules in timelapse images but also perform multiple downstream analyses.

QuantiTrack addresses this need through a graphical user interface, which guides users through each step of the workflow. QuantiTrack is accompanied by a detailed User Guide that walks the user through the software set up and analysis, so that users without prior MATLAB experience can perform and evaluate their analysis independently. The data are stored in a standardized table format on which all the included modules can be run, eliminating the need to reformat the data between steps. The GUI allows the user to load a movie, define regions of interest, detect and track particles, and perform quality control **(Figures 1 and 2)**. QuantiTrack also allows users to draw multiple ROIs to track single molecules in the context of cellular features that are visualized using a different fluorescently labeled molecule: for example, an array of binding sites for a TF^24,47,48^, chromatin and chromatin marks^34,49,50^, or transcription sites^51^. Masks generated in ImageJ or Metamorph can also be imported directly into QuantiTrack (see User Guide).

### Detection and tracking fidelity

Robust particle tracking is essential for robust SMT analysis. Given the volume of data that an SMT experiment can generate, manual inspection of every trajectory is generally not feasible (although this has been done in at least one previous study^52^). To simplify the quality control process, QuantiTrack includes a dedicated Preprocessing panel **(Figure 2)** that generates QC metrics.

We simulated SMT movies of normal diffusion to demonstrate how to use these QC metrics to optimize the detection threshold **(Figure 2, Supplementary Figures 1–2)**. The Kolmogorov-Smirnov (KS) statistic, which measures the separation between the signal and local background intensity distributions can be used to determine the detection threshold at which the two distributions are sufficiently separated **(Figure 2c)**. In practice, this detection threshold can be selected by calculating the KS statistic for a representative set of movies and identifying the inflection point of the KS statistic vs detection threshold curve **(Figure 2n–o and Supplementary Figure 2)**.

By varying the signal intensity, we showed that detection fidelity depends strongly on particle intensity **(Figure 2 and Supplementary Figure 2)**. Signal intensity can be increased by increasing the excitation laser power, with the tradeoff that higher excitation powers also increase photobleaching, thereby reducing track lengths and potentially inducing phototoxicity. By varying the diffusion coefficient, we also demonstrated that motion blur degrades detection fidelity for rapidly diffusing species **(Figure 3)**. Stroboscopic excitation with very short excitation pulses (and high laser powers) can mitigate this to some extent but requires a more complicated optical setup.

Tracking fidelity can be optimized by appropriately selecting the maximum jump distance allowed for each particle such that the probability of observing jump distances longer than the maximum jump is low **(Figure 2e, Supplementary Figure 3)**. We showed that tracking fidelity worsens for high diffusion coefficients largely resulting from individual long trajectories being fragmented into multiple short ones (i.e. reduction in completeness; **Figure 4)**. A maximum jump that is too small prematurely truncates trajectories and biases them towards shorter jumps. On the other hand, very long maximum jump values can result in incorrect linkage of one molecule to another neighboring molecule.

The consequences of FN and FP detections depend on the downstream analysis being performed. FNs result in long trajectories being fragmented into multiple shorter trajectories. Measurements involving successive displacement events that are pooled across all trajectories (such as Spot-On and turning angle distributions) are less susceptible to FN errors, as long as the FN are not associated with a particular mobility state. However, FP detections that result in mistracking still affect these metrics since they introduce spurious displacements. On the other hand, measurements that depend on track length (such as dwell time distributions and MSD analysis) are particularly sensitive to FN detections and resulting trajectory fragmentation. Thus, detection and tracking parameters should also be selected with the down-stream analysis in mind.

Appropriate labeling density also depends on the diffusion coefficient of the protein of interest, and over-crowding can be assessed from the distribution of second nearest neighbor distances provided in the Preprocessing window **(Figure 2f)**. While a labeling density of ∼30 particles/frame is acceptable when tracking a particle diffusing at 1 μm^2^/s, anything over 10 particles/frame significantly degrades the tracking fidelity of a particle diffusing at 5 μm^2^/s **(Supplementary Figure 4)**. These numbers are not universal benchmarks but depend on the imaging and experimental parameters (largest diffusion coefficient, exposure time and imaging interval used in the microscopy setup, particle intensity, and fluorophore choice). While we have focused on SMT with conventional fluorophores, the use of photoactivatable fluorophores can mitigate labeling density issues by activating a single molecule at a time, but this requires additional lasers. We recommend performing initial parameter optimization in the Preprocessing window using three to five representative movies.

These recommendations do not imply that tracking parameters can be adjusted to fix issues with labeling density, laser power, or acquisition parameters. For example, if the signal intensity is too low (because of too short exposure times, low laser powers, or high background), raising the detection threshold will result in the detection of only the brightest particles. Similarly, if the labelling is very dense, reducing the maximum jump could give the appearance of reducing mistracking, but would result in artificially constrained tracks that would bias measurements of spatial mobility such as jump distances or MSD. These issues can be mitigated in the early stages of designing SMT experiments by inspecting the movies of tracks overlaid on the raw images, using the ‘Make Movie of Tracks’ button in the preprocessing step **(Figure 2)** and revisiting the choice of imaging, labeling and other parameters.

### Complementary analyses for resolving mobility features

The Post-processing panel allows the user to select from multiple analysis options that can be run on a set of trajectories **(Figure 1, panel 5)**. These analyses quantify the spatial and temporal dynamics of the protein of interest, provide measures of uncertainty, and generate publication-quality figures and movies.

The MSD can be fit to a power-law model to estimate the diffusion coefficient and scaling exponent. By analyzing simulated trajectories, we showed that diffusion coefficients up to 5 μm^2^/s could be reliably recovered at a frame interval of 12 ms **(Figure 4f–i)**. This upper limit depends on the specific simulation parameters and should not be interpreted as a universal limit of SMT.

While MSD analysis can be used to quantify the motion of trajectories drawn from a single mobility state, it cannot resolve the mobility parameters of a heterogenous collection of trajectories drawn from multiple mobility states. To address this, we benchmarked three complementary spatial mobility analyses (Spot-On, RL, and pEMv2), against simulations of two-state systems with varying proportions of a fast (D_fast_ = 2.5 μm^2^/s) and slow (D_slow_ = 0.1 μm^2^/s) diffusive state **(Figure 5c–f, Supplementary Figure 5a–c)**.

Spot-On jointly fits a two- or three-state model to jump distance distributions across multiple lag times, providing population-level estimates of diffusion coefficients and population fractions. Spot-On pools displacements across all trajectories, and thus, it does not assign states to individual trajectories and does not allow for transitions between states.

QuantiTrack also implements the full kinetic model developed by Mazza et al.^15^. Unlike Spot-On, this explicitly models transitions between different states and estimates transition rates, although at a computational cost. We refer the reader to the User Guide for more information on this.

RL analysis iteratively fits the jump distance distribution at a single lag time to estimate the distribution of MSDs. Individual tracks can be assigned to distinct mobility groups, allowing the user to calculate other parameters (for example MSD exponents, diffusion coefficients, turning angle anisotropy) for the trajectories assigned to each group.

pEMv2 provides the most granular information since state assignment is performed on sub-tracks. While pEMv2 is more computationally expensive than RL and Spot-On, it does not require the user to specify the number of states or type of motion and can identify state transitions between consecutive sub-tracks within a single trajectory.

Both RL and pEMv2 were applied to the simulated two-state system, and each recovered the population fractions and mobility parameters of the two simulated states **(Figure 5g–n, Supplementary Figures 5d–l and 6)**.

We note that in addition to the results reported here, Spot-On^5^, RL analysis^9,31^, and pEMv2^32^ have been rigorously benchmarked in their respective source publications. Together, these approaches provide complementary information with different levels of detail, each with their own strengths and limitations.

### Turning angle analysis provides an exploratory measure of directional motion

The directional persistence of consecutive displacements can provide information about how particles explore their local environment^16,17,35^. Normal, sub-, and super-diffusion were used to illustrate how turning angle distributions and anisotropy coefficients distinguish between these different types of motion. Our simulations further showed that localization error, arising from motion blur, can bias the turning angle distributions towards forward jumps **(Figure 6f–k)**. We must emphasize that the turning angle analysis is the most exploratory component of QuantiTrack. Further modeling and simulations will be necessary to draw quantitative conclusions from this analysis.

When either one of the two displacement vectors making up the turning angle are comparable to (or smaller than) the localization precision, the calculated turning angle is likely dominated by the localization error. QuantiTrack thus includes an option to exclude, from the turning angle analysis, any displacement pair where even one of the two displacements is smaller than the localization precision. For a more rigorous quantification, we suggest running vbSPT to identify and remove the bound segments within all the trajectories and run the turning angle analysis only on the ‘free’ segments^17,34^.

### QuantiTrack reveals hormone-dependent changes in GR dynamics

To demonstrate the utility of QuantiTrack, we applied it to quantify changes in GR dynamics in response to hormone treatment and washout. This is a system that our lab has studied extensively using FRAP and biochemical techniques. Using a combination of fast and slow SMT, we showed that throughout the nucleus, GR dynamics rapidly respond to changes in cellular hormone concentration. Hormone washout resulted in a reduction in the GR bound fraction **(Figure 7h)**, a significant decrease in GR dwell times **(Figure 8d)** and the occupancy of GR in the lowest mobility state at the level of sub-tracks (which is correlated with active GR; **Figure 8f**). These changes in GR binding and dynamics are subsequently reflected in the downregulation of GR target genes **(Figure 7c)**.

Although the biological focus here has been on studying TF dynamics, QuantiTrack is compatible with any SMT dataset. The only difference is that instead of histones, a different photobleaching probe would need to be selected for the quantification of lifetimes. Alternatively, this analysis can be performed without photobleaching correction, but in this case any comparisons would be non-quantitative.

## LIMITATIONS

While developing QuantiTrack, we focused on creating a tool that provides a broad range of tracking and analysis capabilities within a single program. That said, several other analyses like vbSPT^53^, TrackIt^19^, state-array SPT^6^, among others have not been included in QuantiTrack, but can provide additional or complementary information. These tools are freely available and can be applied to QuantiTrack trajectories after appropriate reformatting.

We chose to develop this program in MATLAB since many MATLAB-based image analysis tools exist, which would lower the barrier-to-entry for a new user. However, MATLAB being a proprietary software, we do appreciate that not every user may have access to it.

An astute reader would have noticed that we use the term ‘bound’ and ‘stopped’ molecules interchangeably. We must emphasize that at this time, SMT does not have the resolution to identify site-specific binding events^4^. As a result, anything we call ‘bound’ refers to a spot that is stopped.

Finally, results obtained from QuantiTrack depend on the acquisition and analysis parameters. Different models and parameter choices could yield different estimates of mobility parameters. Thus, while absolute numbers may change, relative comparisons between a control and test dataset are likely to persist when the data are acquired and analyzed under identical conditions. The simulations represent idealized datasets and cannot capture the full kinetic complexity of experimental biological data. Our application of QuantiTrack to experimental GR data therefore serves to complement the simulation benchmarks by applying the analyses to a more heterogenous dataset.

## METHODS

### Cell culture

MCF-7 cells were grown in Dulbecco’s Modified Eagle Medium (DMEM, Thermo Fisher Scientific # 10313039) supplemented with 10% fetal bovine serum (FBS, Gemini Bio, #900-308-500), 2 mM L-glutamine (Thermo Fisher Scientific #25030081), 1% MEM non-essential amino acids (Thermo Fisher Scientific #11140050), 1 mM sodium pyruvate (Thermo Fisher Scientific #11360070), and 50 U/mL penicillin and 50 μg/mL streptomycin (Thermo Fisher Scientific #15070063).

### Generation of H2B-Halo and Halo-GR cell lines

To insert the HaloTag at the N-terminus of the *NR3C1* gene (NM_000176.3), which encodes the glucocorticoid receptor, gRNAs were cloned into the lentiCRISPR v2 (Addgene #52961) plasmid, which contains the Cas9 and puromycin resistance cassette **(Supplementary Figure 8a)**. This cell line was validated by Western blot **(Supplementary Figure 8b)**. GR was detected by chemiluminescence using the GR G-5 antibody (Santa Cruz Biotechnology # sc-393232).

To insert the HaloTag cassette at the C-terminus of the human *HIST1H2BG* gene (NM_003518), we designed a donor plasmid containing the HaloTag sequence flanked upstream by a homology arm (∼800 bp) and 48 bp TEV linker, and downstream by a stop codon and a homology arm (∼800 bp) **(Supplementary Figure 8c)**. These reagents have been previously validated for human cell lines^54^.

5×10^6^ MCF-7 cells at ∼70% confluency were trypsinized and resuspended in 100 μL of DMEM without FBS or antibiotics and electroporated with the respective donors and two gRNA-containing lentiCRISPR v2 plasmids. We used 2 gRNAs to improve the CRISPR efficiency. For the electroporation we prepared 5 million cells in 100 μl of media (without FBS or antibiotics) with 5 μg of each plasmid. We used the BTX T820 Electro Square Porator (Harvard Apparatus, Holliston, MA, USA) and applied 3x 10 ms pulses of 120 V. After electroporation, the cells were resuspended in complete fresh media. Since the cells transfected with the lentiCRISPR v2 plasmid acquire temporary resistance to puromycin, the electroporated cells were grown in the presence of 1 μg/mL puromycin for two days. The surviving cells were washed three times with PBS and grown in medium without antibiotics until they reached 10×10^6^ cells.

Cells were then incubated with 5 nM Halo-TMR ligand (Promega #G8251) for 20 min, washed three times, and the washed one more time after a 10 min incubation period. HaloTag expressing cells were then selected by fluorescence-activated cell sorting (FACS) and single positive clones were grown in 96 well plates. Several clones were selected and expanded further. Finally, we selected and validated one clone for each of the cell lines for this study.

### RNA-qPCR experiments

For RNA-qPCR experiments, MCF-7 cells were plated in 6 well plates in medium containing charcoal-stripped FBS for 24 hours. In the treatment protocol, cells were treated with 1.2 μM Hcort (Sigma Aldrich #H4001-1G) for 20 min or 60 min. In the washout protocol, MCF-7 cells were treated with 1.2 μM Hcort for 20 min and then washed three times with conditioned medium. RNA was collected at 20 min and 60 min using the PureLink RNA Mini Kit (ThermoFisher Scientific #12183025) following the manufacturer’s protocol. For the control samples, cells were treated with 0.1% EtOH for 60 min, and this serves as our baseline.

### Optical-dynamic simulations

Optical-dynamic simulations were performed using the sptPALMsim^29^ python package. The particle trajectories are drawn from fractional Brownian motion (fBM) with the generalized diffusion coefficient (D_α_) and Hurst parameter (H = α/2). The trajectories simulated by sptPALMsim span a 3D volume of ∼ 40 × 40 × 10 μm^3^. Of this, the central ∼ 13.3 × 13.3 × 4 μm^3^ is rendered by the optical and camera model. To generate the movie, each trajectory is convolved with the 3D PSF specified by the input parameters. When motion blur is enabled, each frame is divided into 60 micro frames, and the particle coordinates are integrated over the full frame interval.

Unless otherwise specified in the figures, the default simulation parameters were as follows:

| Parameter | Value |
| --- | --- |
| --frame_interval | 0.012 s |
| --pulse_width | 0.010 s |
| --bleach_rate | $0.2 \text{ s}^{-1}$ |
| --wavelength | $0.67 \mu\text{m}$ |
| --na | 1.45 |
| --ref_index | 1.515 |
| --pixel_size | $0.104 \mu\text{m}/\text{px}$ |
| --fov_size | 128 px |
| --spatial_bin_rate | 8 |
| --offset | 425 ADU |
| --gain | $75 \text{ e}^-/\text{ADU}$ |
| --read_noise | $3 \text{ e}^-$ |
| --intensity | 150 photons/frame |
| --n_tracks | 1500 |
| --n_frames | 300 |
| --n_reps | 10 |
| --z_half_range | $2.0 \mu\text{m}$ |
| --H | 0.5 |

To construct the ground truth trajectories, the lateral bounds of the trajectories were defined according to the 128 × 128 px (13.3 μm × 13.3 μm) ROI and the axial range was set to ± 0.35 μm (total range of 0.7 μm).

The ground truth trajectories were clipped whenever a localization fell beyond these xyz limits.

For the density sweep **(Supplementary Figure 4)**, - -n_tracks was set to 1500, 3000, 5000, 10,000, 20,000, corresponding to 10, 20, 30, 50, 90 visible particles per frame, respectively. To reduce computational cost, for the -- n_tracks = 10,000 and 20,000, the simulations were limited to 100 frames. Except for the slow/fast mixture simulations, 10 movies were simulated for each parameter set.

For each fast/slow mixture **(Figure 5, Supplementary Figures 5 and 6)**, 120 movies were simulated per dataset.

### Detection fidelity metrics

Pairwise Euclidean distances between the QuantiTrack and ground truth (GT) localizations were calculated using the MATLAB function pdist2. Distances exceeding 2 pixels (208 nm) were set to Inf. The one-to-one assignments were then performed using matchpairs to obtain the true positives (TP). Unmatched QuantiTrack and GT localizations were counted as false positives (FP) and false negatives (FN) respectively.

The detection fidelity was calculated using the Jaccard similarity coefficient (JSC)^30^

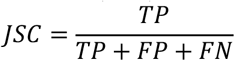

### Tracking fidelity metrics

The overlap matrix *O* was constructed using accumarray such that *O*(*k, j*) is the number of frames in which the GT trajectory *X*_*k*_ and QuantiTrack trajectory *Y*_*j*_ matched. The one-to-one track assignments between the GT and QuantiTrack tracks were made using *matc*h*pairs*(−*O*, 0). A pair was counted as a true positive track (TPt) only if more than 50% of the QuantiTrack track’s localizations matched those of the assigned GT track^30^. Unmatched QuantiTrack and GT trajectories were counted as false positive (FPt) and false negative (FNt) tracks respectively.

The track-level JSC was then computed as

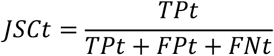

The purity (*P*) is calculated as the fraction of QuantiTrack localizations that map on to their matched GT trajectory:

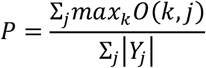

The completeness *C* is calculated as the fraction of GT localizations that map on to their respective QuantiTrack trajectory:

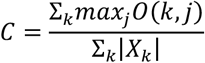

### Single molecule tracking sample preparation

100,000 MCF-7 cells expressing Halo-GR or H2B-Halo were plated in 2 well chamber slides (Cellvis #C2-1.5H-N) in charcoal-stripped FBS supplemented medium for 24 hrs. On the day of imaging, cells were incubated with 50 pM JF549 (Promega # GA1110; for the slow SMT experiments) or 100 pM JFX650 (Promega #HT1070; fast SMT experiments) for 15 min. The samples were washed three times with phenol red-free charcoal-stripped FBS containing medium. The samples were returned to the incubator for 10 min and washed one more time.

After the dye labeling step was complete, the samples were treated with 1.2 μM Hcort for 20 min. For the washout condition, the sample was washed three times with phenol red-free charcoal-stripped FBS medium, while nothing further was done to the samples in the treatment protocol. The chamber slides were then allowed to equilibrate on the microscope for 10 min before collecting movies. Data were collected between 30 and 60 min after initial hormone addition.

### Microscopy

All the data reported in this study were collected on a custom-built highly inclined and laminated optical sheet (HILO) microscope at the Optical Microscopy Core of the NCI. This system has been described and characterized in detail in a previous publication^55^. Briefly, this microscope is built on the body of an Olympus IX81 inverted microscope and is equipped with a 150×, 1.45 NA objective (Olympus Scientific Solutions, Waltham, MA, USA). This system has a 561 nm laser (iFLEX-Mustang, Excelitas Technologies Corp., Waltham, MA, USA), a 647 nm laser (OBIS 647 LX, Coherent Inc, Santa Clara, CA, USA), an acousto-optical tunable filter (AOTFnC-400.650, AA Optoelectronic, Orsay, France), and an EM-CCD camera (Evolve 512, Photometrics, Tucson, AZ, USA). For live cell-imaging, the microscope has a stage-top incubator with 5% CO_2_ control (Okolab, Pozzuoli NA, Italy).

For fast SMT experiments, images were collected every 12 ms under continuous illumination by the 647 nm laser **(Figure 7d)**. For slow SMT experiments, movies were collected under stroboscopic illumination by the 561 nm laser with 10 ms exposures at a frame rate of 5 Hz (200 ms intervals, with a 190 ms dark period between consecutive illuminations) **(Figure 8a)**. Both laser powers were set to 0.96 mW measured at the objective back aperture. The pixel size for this setup is 104 nm/pixel.

### Tracking

Image stacks were filtered using a top-hat, Wiener, and bandpass filter. The nuclear periphery was defined using a hand-drawn region of interest based on the sum projection of the stack. QuantiTrack can also import masks from ImageJ and Metamorph (see User Guide). The detection threshold parameter was determined as the inflection point of the KS statistic vs detection threshold curve **(Figure 7d, S9)**. Particles were localized by fitting a Gaussian point spread function. The fitted model assumes a fixed Gaussian width, which we found to provide more robust position information compared to versions where the width was allowed to vary. Detected particles were connected into tracks based on a nearest neighbor algorithm. For the fast SMT data, the maximum jump was set at the point where the probability of jumps longer than MJ was negligible. MJ = 6 px (624 nm) and 8 px (832 nm) for GR + Hcort treatment and washout respectively. For the slow GR SMT data, we focused on the bound molecules and hence set MJ = 4 px (416 nm) as has been used previously^7,18^.

The simulations were tracked in the same manner as the GR data, with gaps to close = 2 frames, shortest track = 2 frames, and maximum jump as described in the respective figures.

### Quality control

We developed two types of quality control measures that can be applied to each movie while it is being analyzed, or later – on an ensemble of movies. We describe these below:

#### Kolmogorov-Smirnov (KS) statistic

For every detected particle, the particle intensity is calculated as the average intensity over the Airy disk, which has a radius (in pixels), *r*_*Airy*_ given by 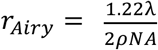, where λ is the wavelength of the emitted light, ρ is the pixel size and *NA* is the numerical aperture of the objective. The local background is calculated as the mean intensity over a donut-shaped region with inner and outer radii (in pixels) calculated as *r*_*inner*_ = *r*_*Airy*_ + 2 and *r*_*outer*_ = *r*_*Airy*_ + 3 respectively **(Figure 2c**, callout).

For a given detection threshold, the KS statistic was calculated as the largest vertical distance between the cumulative distribution function of the particle signal intensity and local background intensity. If *F*_*signal*_(*i*) and *F*_*background*_(*i*) are the CDFs of the signal and background intensities respectively,

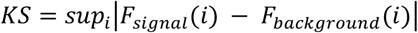

The optimal detection threshold is calculated as the inflection point of the KS statistic vs detection threshold curve **(Figure 2o)**.

#### Signal-to-noise ratio

For a given detection threshold, the estimated signal-to-noise ratio (SNR) for each particle is calculated as

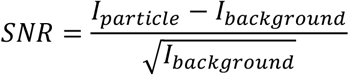

where *I*_*practice*_ and *I*_*background*_ are the particle and local background intensity. A more accurate SNR value could be obtained by adding parameters (gain, offset, read noise, etc.) from the specific camera used to capture the movies. However, the above equation provides an adequate estimate to evaluate particle detection.

#### Maximum jump parameter and jump distance histogram

To capture both fast diffusing and bound trajectories, the maximum jump parameter was calculated as the smallest MJ at which the probability of observing displacements larger than MJ was negligible. For the slow SMT data, we focused on the bound trajectories; hence, MJ was set to 4 px (416 nm).

### Localization precision

The precision with which we estimate the position of each particle is calculated from the 95% confidence intervals of the point spread function (PSF) centroid coordinates (*CI*_*x*_, *CI*_*y*_) obtained from the fit of the spot to a 2D gaussian. Assuming the centroid parameters are normally distributed, the localization precision, σ_*total*_, can be calculated 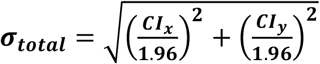 . For each image QuantiTrack reports the median σ_*total*_ from all detected particles.

### Pixel size measurements

Like any quantitative analysis of imaging data, QuantiTrack relies on a conversion from pixels to physical distance. It is therefore imperative to accurately determine the pixel size, *px*_*Im*_, within your images to ensure that downstream estimates of displacements and diffusion coefficients reflect real values. In commercial microscope systems, this will often be explicitly specified or can easily be calculated as 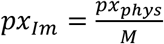, where *px*_*phys*_ is the physical size of a pixel on the CCD (or sCMOS) chip, and *M* is the magnification of the objective lens.

If using a custom-built microscope or if the physical size of the camera’s pixel is unknown, we recommend measuring the pixel size before running an experiment. This can be achieved with a motorized stage using beads mounted on a slide, such as the TetraSpeck kit (ThermoFisher #T14792), by moving the stage a specified distance and dividing by the number of pixels the bead moved in the images. Microscope control software, such as the open-source Micromanager 2.0^56^, also usually contains pixel calibration tools to obtain more precise measurements of sizes.

### Dwell time distribution

To identify bound segments within a track, the user must determine the distance a stably bound molecule moves over consecutive frames (*r*_*min*_). Since transcription factors interact with chromatin, we used histone H2B as our reference molecule, and the parameter *r*_*min*_ is calculated as the 99^th^ percentile of the H2B frame-to-frame displacements. Freely diffusing molecules can move less than this threshold distance in one frame, but the probability that a freely diffusing molecule with diffusion coefficient *D*_*free*_, moves a distance *r*_*min*_ over each of *N*_*min*_ frames in time Δt is given by

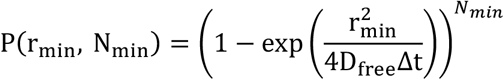

We used this equation to determine the number of frames *N*_*min*_ over which the probability of classifying a diffusing molecule as bound is less than 1%. We then determined the distance *r*_*max*_, which is the 99^th^ percentile of the H2B displacement histogram over *N*_*min*_ frames, and further eliminated track segments where the TF displacement was larger than *r*_*max*_ over *N*_*min*_ frames. A survival distribution was then compiled from all the bound segments.

### Photobleaching correction

The histone H2B survival distribution was used to measure the photobleaching rate and correct for photo-bleaching using our previously described method^18^. We generated the survival distribution for H2B as described in the section ‘Dwell time distribution’. We fit this survival distribution to a triple exponential function and extracted the slowest exponential decay rate as the photo-bleaching rate *k*_*PB*_ . The raw survival distribution *S*_*raw*_(t) is then corrected for photobleaching as follows, where *t*_0_ is the first point of the survival distribution:

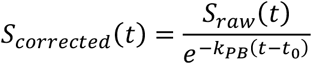

Finally, we normalize the survival distribution to the bound fraction such that:

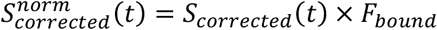

### RL analysis

The self-part of the jump distance histogram or van Hove correlation (vHc) was calculated as *G*_*s*_(*r, τ*) = *A*_*s*_⟨*δ*(*r*_*i*_ − |*r*_*i*_(t + *τ*) − *r*_*i*_(*t*)|⟩, where ***r***_***i***_ is the position of the i^th^ particle and *A*_*s*_ = ∫ *d*^2^ *rG*_*s*_(*r, τ*) is a normalization constant. As done previously^7,9^, we assumed that the vHc can be expanded as a superposition of Gaussian functions 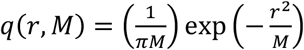 such that *G*_*s*_ (*r, τ*) = ∫ *P* (*M, τ*)*q*(*r, M*)*dM*, where *M* is the mean-squared displacement and *P*(*M*) is the probability density of the mean-squared displacements of the ensemble of particles. We then used an iterative scheme developed by Richardson and Lucy and recently adapted to nucleosomes^9^ and other transcriptional proteins^7,25^. From an initial distribution 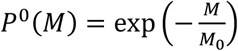, the distribution in the (*n* + 1)^th^ iteration, *P*^*n*+1^, can be written as

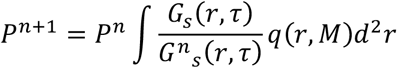

Since *P*^*n*+1^ is a probability density, we impose the constraint *P*^n+1^ > 0 and that *P*^*n*+1^ be normalized.

To classify into different mobility groups, we first identified the local minima in the inferred MSD distributions **(Figure 8i and Supplementary Figure 5g–i)**. We used these local minima to bin the MSD distribution into different mobility groups. Each track was assigned to the bin corresponding to its MSD at the specified time lag as previously described^7,25^.

### Perturbation expectation maximization v2 analysis

Perturbation expectation maximization v2^32^ was run as described previously^7^. Briefly, all tracks were divided into 7-frame sub-tracks, thus retaining only those tracks that are at least 7 frames long. We chose a split length of 7 frames to minimize transitions between different mobility states within a single sub-track. Different split lengths have been tested previously, yielding similar results^24^. The system was allowed to explore between 1 and 15 states with 50 reinitializations and 200 perturbations. Up to 10,000 iterations were allowed and the convergence criteria for the log-likelihood function was set at 10^-7^. Convergence of pEMv2 was verified by running each dataset five times, and the run with the lowest BIC was selected for further analysis.

The output from pEMv2 is the optimal number of states and a posterior probability for each sub-track. We assigned the sub-tracks to the states for which they had the highest posterior probability **(Supplementary Figure 6a)**. For MSD visualization and population fraction calculations, we only considered those sub-tracks for which the difference between the two highest posterior probabilities was greater than 0.2. This was done to ensure that sub-tracks with similar posterior probabilities for multiple states do not contaminate the data.

### Turning angle analysis

Angles formed by 3 successive localizations were calculated for all tracks and plotted as a polar histogram. As previously described by Izeddin, et al.^16^, we calculated the anisotropy coefficient (AC) as the log_2_ ratio of the probability of forward displacements [*prob*(−30° ≤ *θ* ≤ 30°)] to that of backward displacements [*prob*(150° ≤ *θ* ≤ 210°)] as follows **(Figure 6e)**:

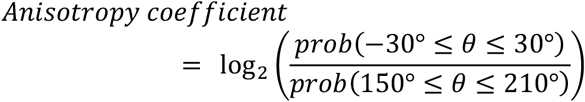

### Bound fraction by Spot-On

Spot-On^5^ was run with the indicated time lags with the following parameters:

TimeGap = 12; dZ = 0.7; GapsAllowed = 1; TimePoints = 7; BinWidth = 0.01; UseEntireTraj = 1; UseWeights = 1; MaxJump = 0.7 (GR + Hcort treatment). MaxJump = 1.5 (GR + Hcort washout); ModelFit = 1; DoSingleCellFit = 0; FitIterations = 5; FitLocError = 1; FitLocErrorRange = [0.01, 0.075]; NumberOfStates = 2; D_Free_2State = [0.05, 75]; D_Bound_2State = [1e-5, 0.1]; D_Free1_3State = [0.1, 25]; D_Free2_3State = [0.1, 25]; D_Bound_3State = [1e-5, 0.1].

Error bars represent the standard deviation of 100 bootstrapped samples with replacement.

## ACKNOWLEDGEMENTS

We would like to thank the following people for beta-testing the software and providing useful feedback: Thomas A Johnson, Hannah LaPoint, My Linh Bui, Fadil Iqbal, Ira Kraft (NCI, NIH), Gregory Fettweis (University of Liege), Diego M Presman (University of Buenos Aires, CONICET). We would like to thank Hannah LaPoint for providing input on the CRISPR schematics. We would also like to thank Anders Hansen and colleagues for making their SMT data^17^ freely available and for the MATLAB implementation of Spot-On^5^, and Alec Heckert for sptPALMsim^29^. We acknowledge support from Gianluca Pegoraro and the High Throughput Imaging Facility (HiTIF [CCR, NCI, NIH]).

The contributions of the NIH author(s) were made as part of their official duties as NIH federal employees, are in compliance with agency policy requirements, and are considered works of the United States Government. However, the findings and conclusions presented in this paper are those of the author(s) and do not necessarily reflect the views of the NIH or the U.S. Department of Health and Human Services, nor does mention of trade names, commercial products, or organizations imply endorsement by the U.S. Government.

## AUTHOR CONTRIBUTIONS

**David A Ball** (Conceptualization, Methodology, Software, Validation, Formal Analysis, Writing – Review and Editing, Visualization), **Kaustubh Wagh** (Conceptualization, Methodology, Software, Validation, Formal Analysis, Investigation, Data Curation, Writing – Original Draft, Writing – Review and Editing, Visualization), **Diana A Stavreva** (Methodology, Investigation, Resources, Writing – Review and Editing), **Le Hoang** (Investigation), **R Louis Schiltz** (Investigation), **Raj Chari** (Resources), **Razi Raziuddin** (Investigation), **Davide Mazz**a (Conceptualization, Methodology, Software, Validation, Writing – Review and Editing), **Arpita Upadhyaya** (Methodology, Software, Validation, Resources, Supervision, Funding acquisition, Writing – Review and Editing), **Gordon L Hager** (Resources, Supervision, Project administration, Funding acquisition, Writing – Review and Editing), **Tatiana S Karpova** (Conceptualization, Resources, Supervision, Project administration, Funding acquisition, Writing – Review and Editing).

## DATA AVAILABILITY

QuantiTrack is available on Github: https://github.com/dball07/QuantiTrack

Representative SMT movies and trackTables for the GR data are deposited in Zenodo:

https://doi.org/10.5281/zenodo.22756685

