## Supplementary figures and table for "QuantiTrack: A unified software to study protein dynamics in living cells"

### QUANTITRACK: AN INTEGRATED GUI-BASED SINGLE-MOLECULE TRACKING AND ANALYSIS TOOL

**SUPPLEMENTARY MATERIALS**

**This PDF includes**

- Supplementary Table 1
- Supplementary Figures 1 to 12
- Movies 1 to 6

**Supplementary Table 1. Summary of analyses implemented in QuantiTrack.**

| Analysis | Source Publications | Assumptions | Limitations | Estimated Parameters | Minimum number of tracks |
|---|---|---|---|---|---|
| Dwell-time analysis | • Mazza, et al. Nuc Acids Res 2012<br>• Presman, et al. Methods 2017<br>• Garcia, et al Nuc Acids Res 2021 | • A stable molecule (such as histones for nuclear proteins) is required to measure the photobleaching rate<br>• Max frame-to-frame jump and max jump within $N_{min}$ frames isolates bound segments within tracks | • Parameters ($r_{min}$, $r_{max}$, $N_{min}$) are determined empirically and can affect results | • Photobleaching rate (from a 3-exponential fit to the stable protein survival distributions)<br>• Raw dwell time distributions<br>• BIC-based model fitting (1- or 2-exponential, power-law, and power-law + exponential) | ~500 ("approximately 20 cells yielding a total of several hundred tracks") |
| Spot-On | Hansen, et al. eLife 2018 | • Gaussian distributed displacements<br>• No transitions between states<br>• 2 or 3 states, specified by user<br>• No correlation between successive displacements | • No transitions allowed between states<br>• Not appropriate for directed motion, where successive displacements are correlated<br>• 3-state model may converge to local minimum | • Population fractions<br>• Diffusion coefficients<br>• Localization precision<br>• 95% confidence intervals for each parameter | 3,000 |
| Jump Histogram Full kinetic model | Mazza, et al. Nuc Acids Res 2012 | • Gaussian distributed displacements<br>• Poissonian binding/unbinding model<br>• Assumes a finite focal thickness, and molecules can't re-enter between frames<br>• Localization error modeled | • The model is phenomenological | • Population fractions<br>• Diffusion coefficients<br>• Binding/unbinding rates ($k_{ON}$ and $k_{OFF}$)<br>• Equilibrium bound fraction $C_{eq} = \frac{k_{ON}}{k_{ON}+k_{OFF}}$ | 1,000 (simulations used 1000 particles) |

| | | | | | |
|---|---|---|---|---|---|
| | | as an additive Gaussian term | | | |
| Turning Angle Anisotropy | • Izeddin, et al. eLife 2014<br>• Hansen et al Nat Chem Biol 2020 | • 2D-projected angles are representative of 3D motion<br>• Displacements must be greater than the localization precision (when $loc_{prec}$ filter is enabled) | • Multiple physical models of exploration can produce the same anisotropy signature<br>• Exploratory analysis that needs to be backed by modeling or simulations | • Turning angle distributions<br>• Anisotropy coefficient (AC)<br>• AC over spatial and temporal lags with 95% confidence intervals | 10,000 |
| Perturbation-expectation maximization v2 (pEMv2) | Koo & Mochrie. Phys Rev E 2016 | • Displacements are assumed to follow a stationary Gaussian process<br>• No transitions assumed within a sub-track | • Optimal sub-track length not known a priori<br>• No memory across segments<br>• Computationally expensive compared to RL and Spot-On. | • Posterior probability of state assignment Population fractions<br>• Diffusion coefficients per state<br>• Localization error per state | 1,500-3,000 (depends on model complexity) |
| Richardson-Lucy (RL) | Ashwin, et al. PNAS 2019 | • Gaussian distributed displacements | • Only verified for specific range of track lengths<br>• Calculated for one time lag at a time | • Van Hove correlation and RL model fit<br>• RL deconvolved MSD distribution<br>• Per state MSD with power-law fits | 300-600 |

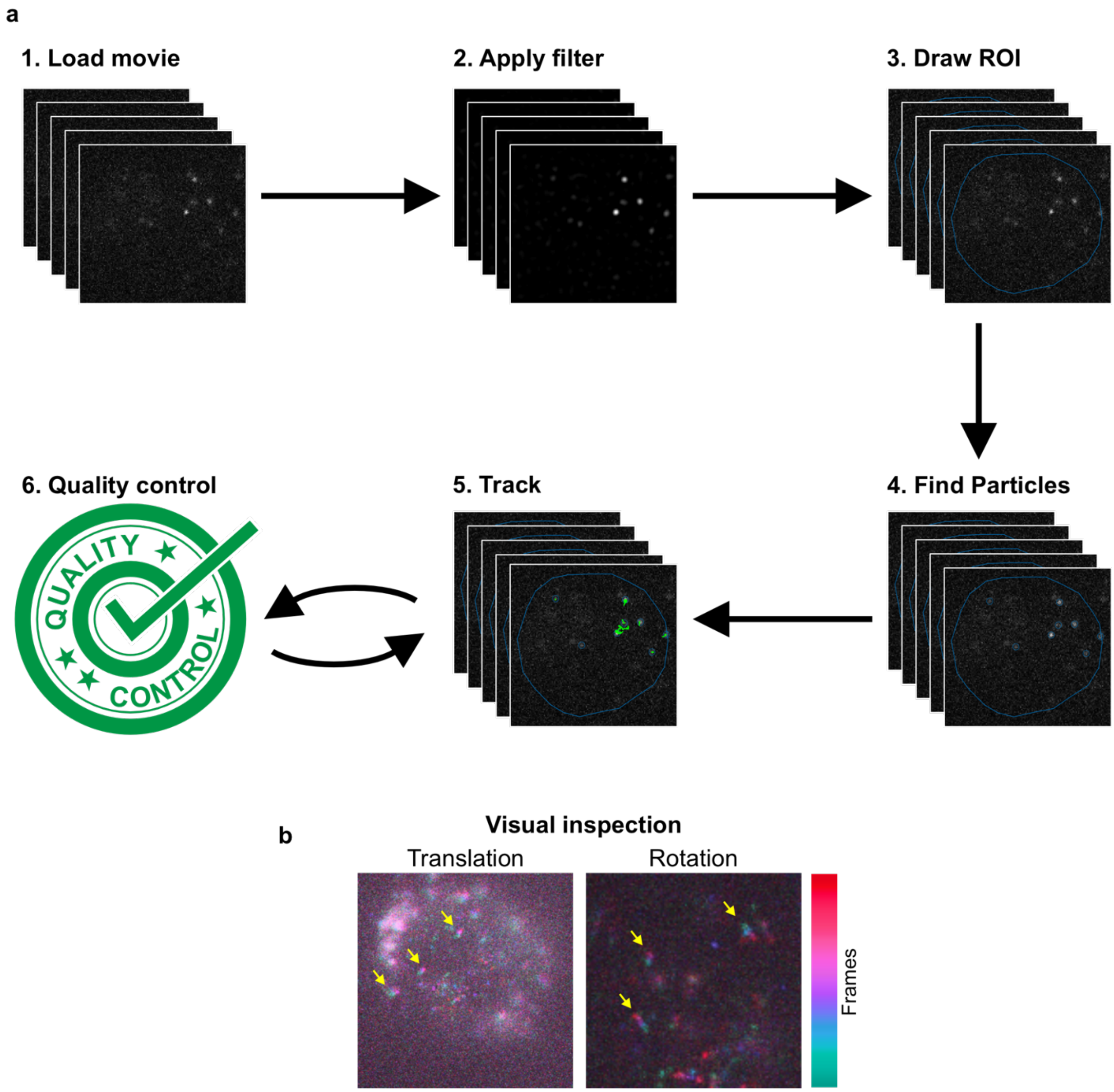


**Supplementary Figure 1. Visual inspection and analysis workflow.**
**(a)** Overview of the QuantiTrack workflow. (1) Load the tif stack into QuantiTrack. (2) Filter the movie using top-hat, Wiener, and Gaussian filters. (3) Draw a region of interest (ROI) to demarcate the nucleus or other features of interest. (4) Find particles. (5) Combine detected particles into tracks. (6) Run the quality control analysis and update tracking or particle detection parameters if needed. **(b)** Temporal projections of movies exhibiting (left) translational drift and (right) rotational drift. The arrows indicate points exhibiting (left) translation and (right) rotation.

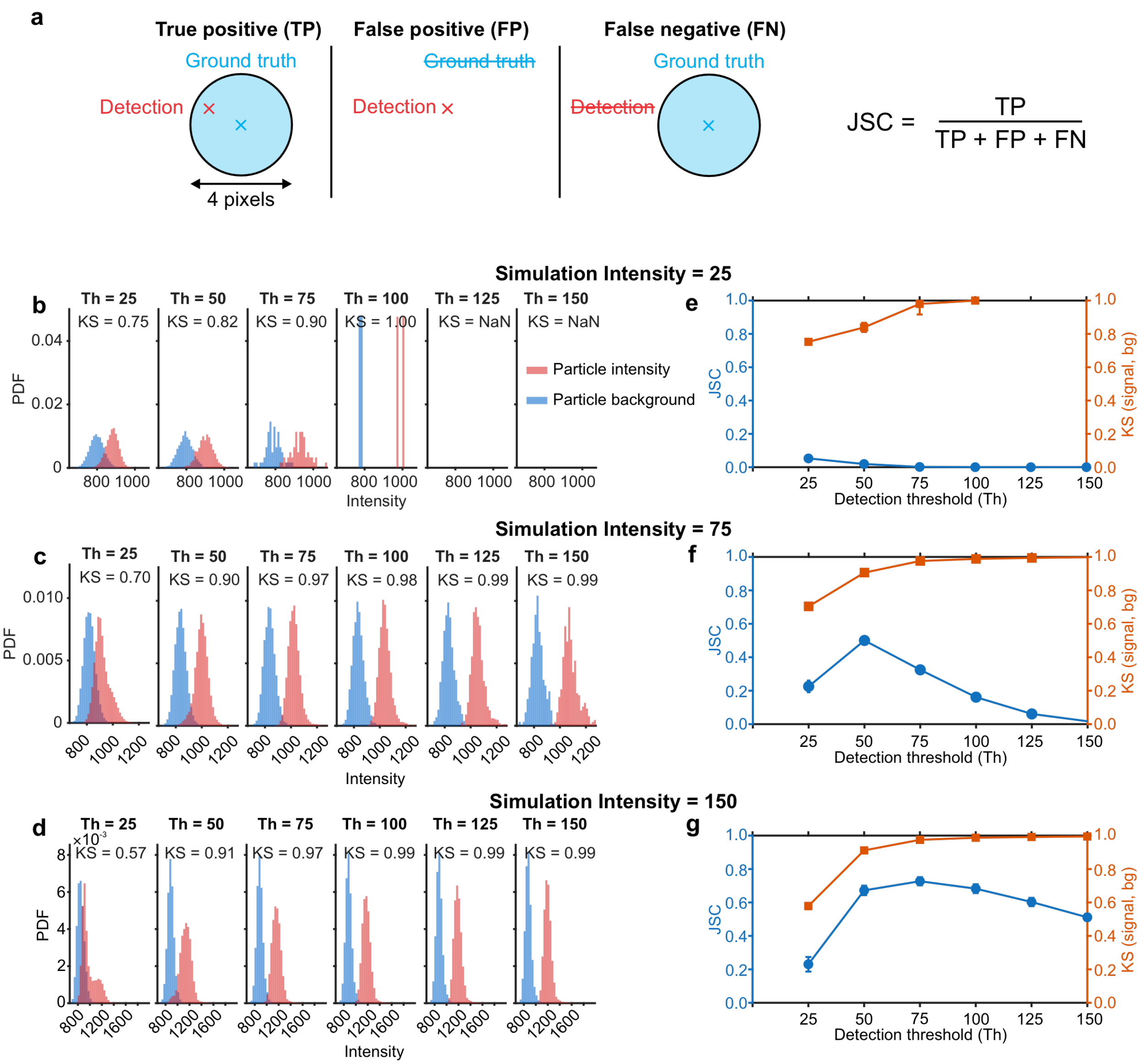


**Supplementary Figure 2. Quality control in QuantiTrack.**
**(a)** Schematic defining true a true positive (TP), false positive (FP), and false negative (FN) detection. The detection fidelity is measured using the Jaccard similarity coefficient (JSC). **(b–d)** Histograms of particle intensity and local background for different detection threshold (Th) values for simulation intensity = (b) 25, (c) 75, (d) 150. Inset text shows the Kolmogorov-Smirnov (KS) statistic measuring the distance between the particle intensity and local background distributions, **(e–g)** The detection JSC and KS statistic as a function of detection threshold for simulation intensity = (e) 25, (f) 75, (g) 150.

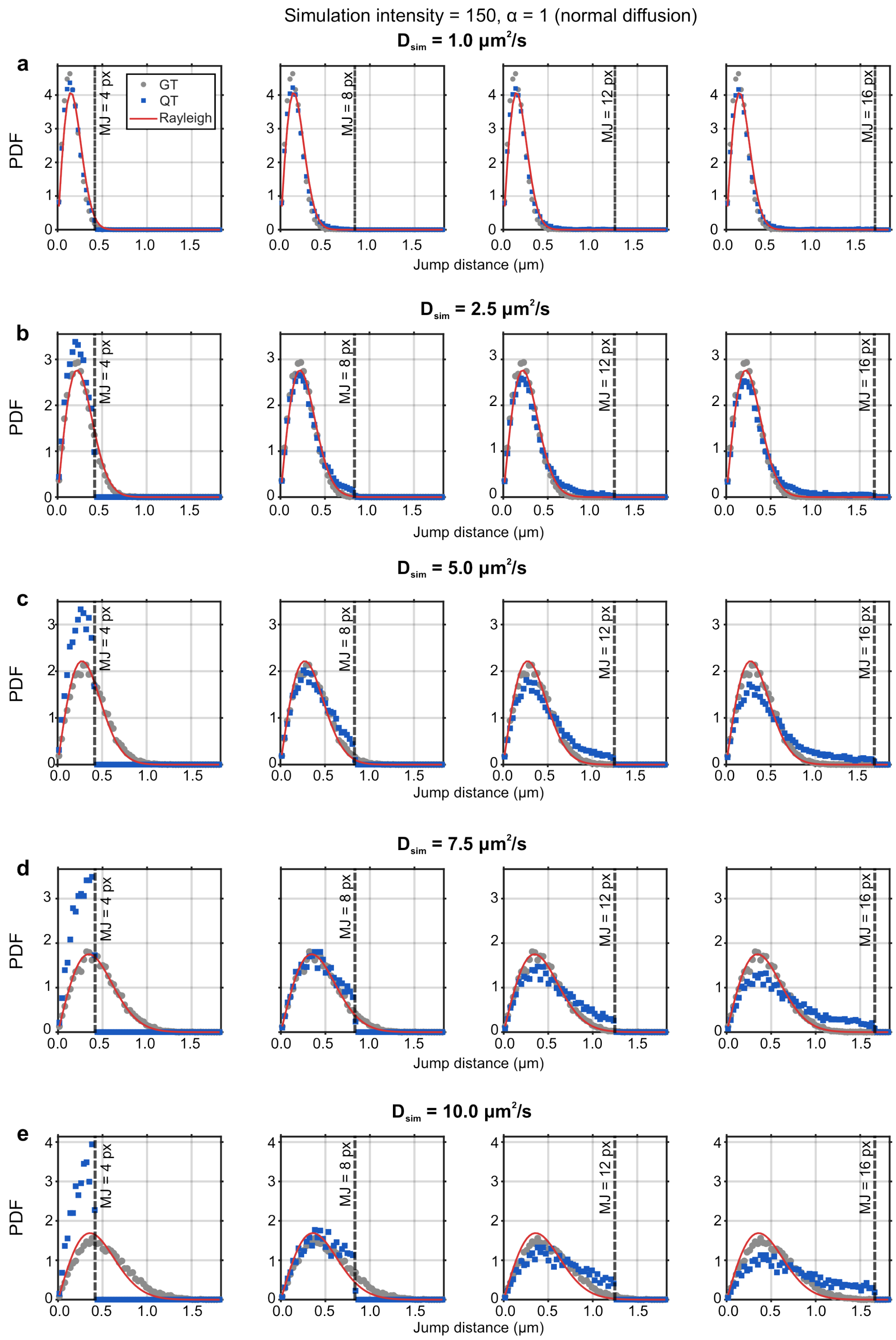


**Supplementary Figure 3. Jump distance histograms.**
Ground truth (GT, gray circles) and QuantiTrack (QT, blue squares) jump distance distributions for normal diffusion with indicated diffusion coefficients and max jump = 4, 8, 12, 16 pixels. The max jump (MJ) distance is indicated with a dashed line. The solid red line is the Rayleigh distribution for each diffusion coefficient.

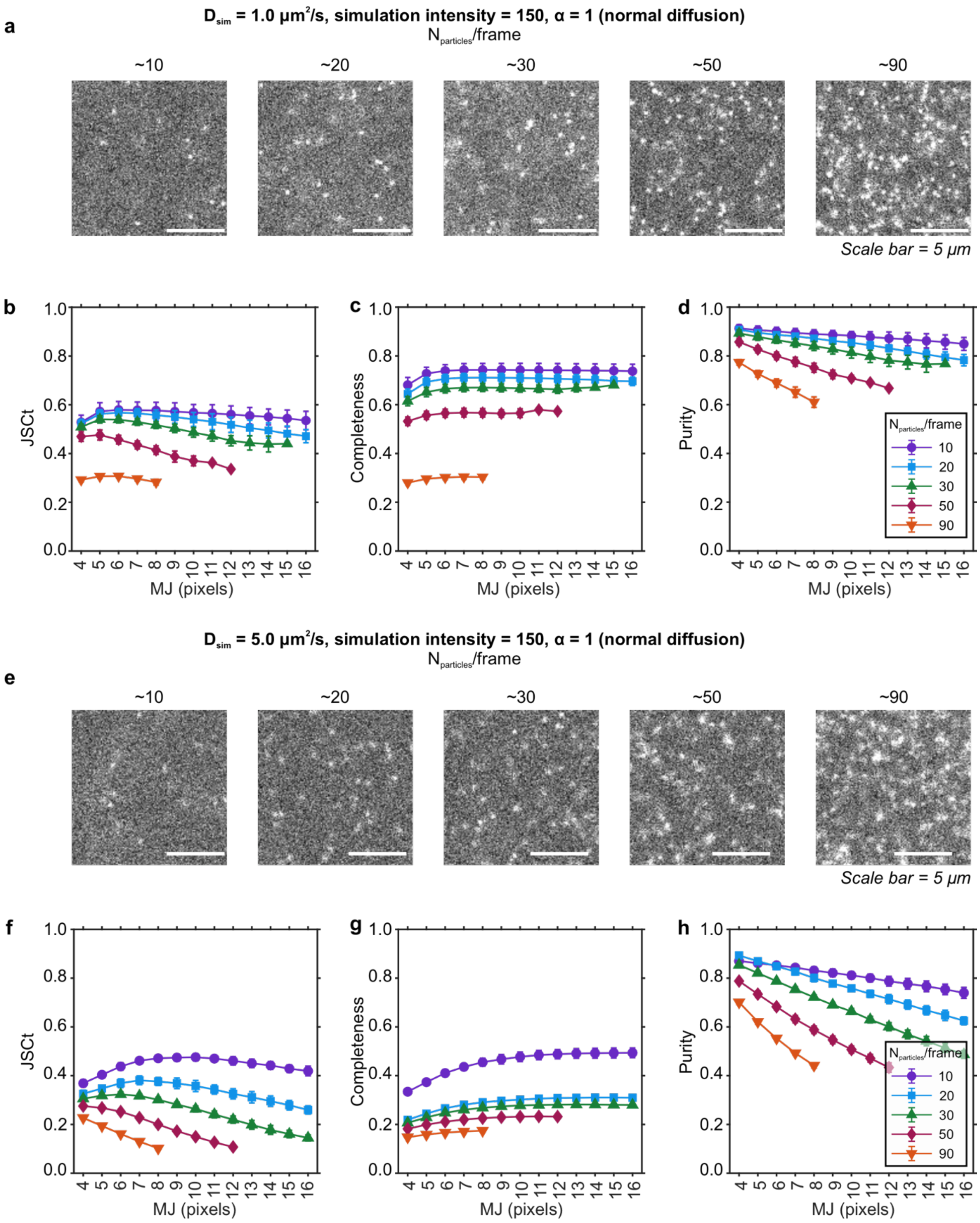


**Supplementary Figure 4. Effect of labeling density on tracking fidelity.**
**(a)** Representative stills from simulations with intensity = 150, D = 1.0 $\mu m^2/s$, $\alpha$ = 1 and increasing labeling density. **(b–d)** Measured (b) track level Jaccard similarity coefficient (JSCt), (c) completeness, and (d) purity as a function of the maximum jump (MJ) parameter for the indicated labeling densities. **(e)** Representative images from simulations with intensity = 150, D = 5.0 $\mu m^2/s$, $\alpha$ = 1 and increasing labeling density. **(f–h)** Measured (f) JSCt, (g) completeness, and (h) purity as a function of max jump for the indicated labeling densities.

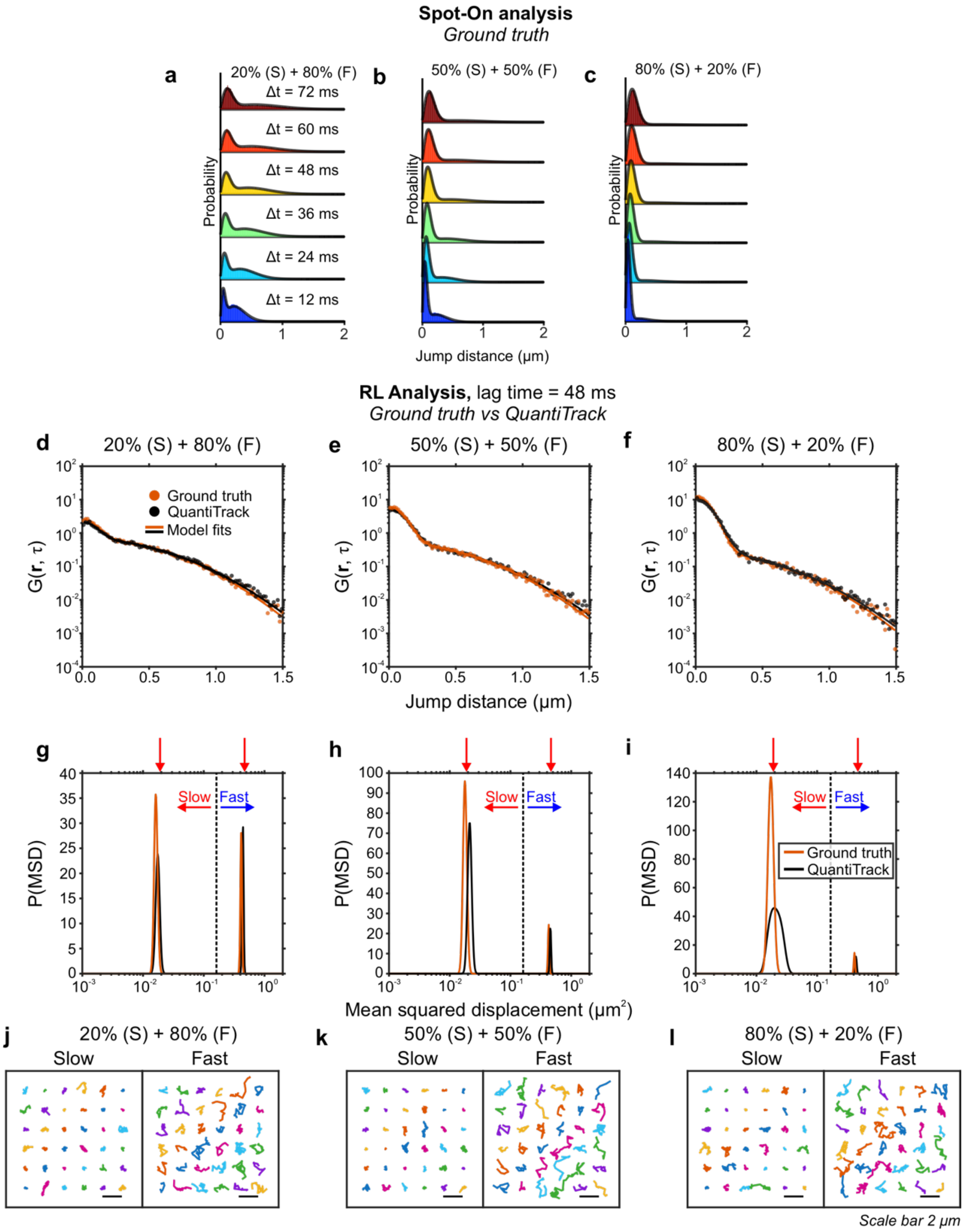


**Supplementary Figure 5. Spot-On and RL analysis of a system with multiple mobility states.**
**(a–c)** Ground truth jump distance distributions and Spot-On two-state model fits (solid lines) for the indicated mixtures of fast and slow states. **(d–f)** van Hove correlation at lag time = 48 ms and Richardson-Lucy deconvolution (RL analysis) model fits (solid lines) for the ground truth (orange) and QuantiTrack data (black) for indicated mixtures of slow and fast states. **(g–i)** Distribution of mean squared displacements (MSD) obtained from the RL analysis for ground truth (orange) and QuantiTrack (black) data for the indicated conditions. Red arrows indicate the true MSD for the slow and fast states (based on the simulation parameters). **(j–l)** Representative trajectories recovered from the RL analysis for the slow and fast states for the indicated mixtures. Scale bar = 2 μm.

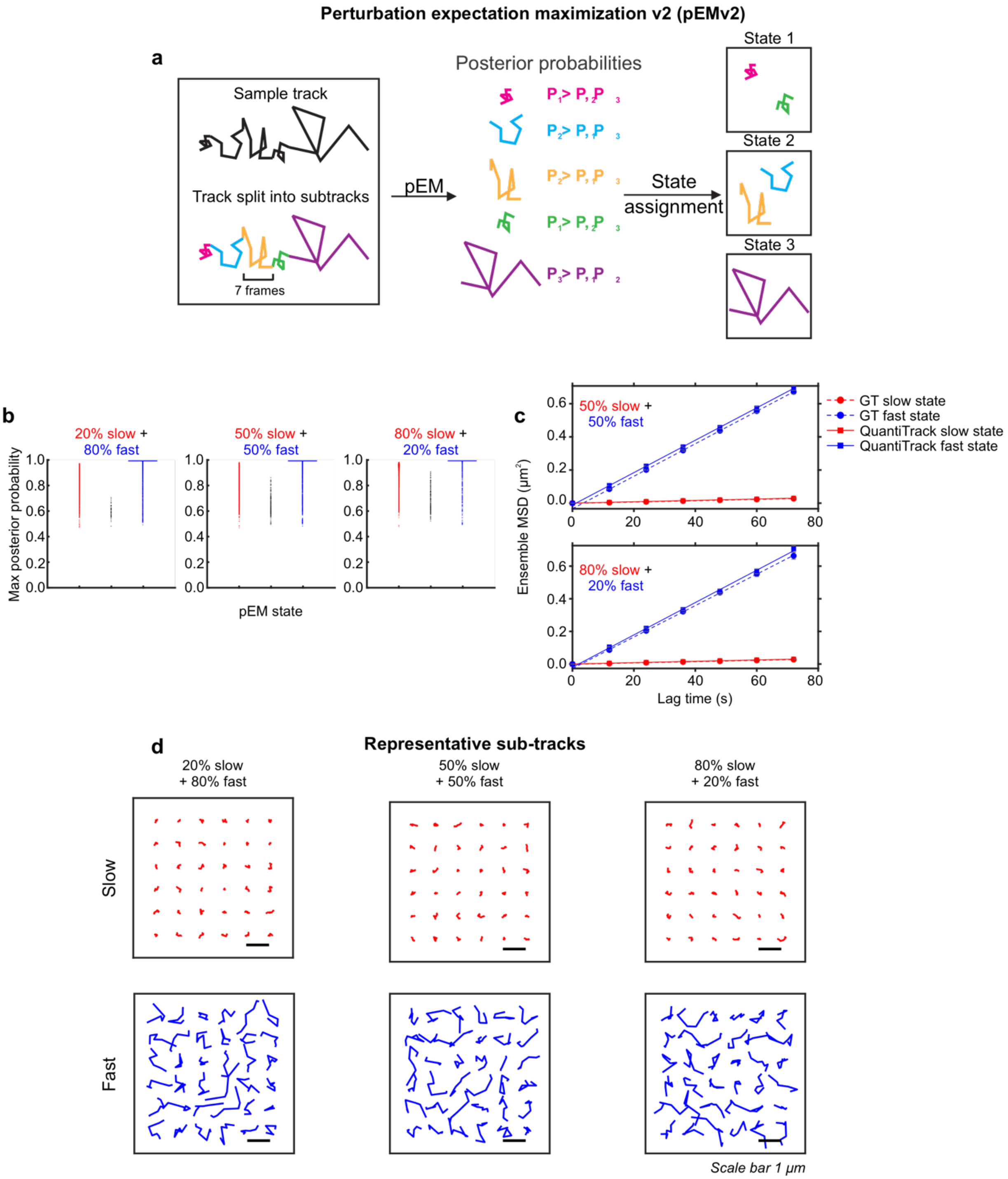


**Supplementary Figure 6. Perturbation expectation maximization v2 (pEMv2) analysis of multiple mobility state systems.**
**(a)** Schematic of the perturbation-expectation maximization (pEMv2) workflow. (Left) Tracks are split into 7-frame sub-tracks. (Center) After running the pEMv2 analysis, each sub-track has a posterior probability of belonging to each mobility state. In this schematic, pEMv2 converged to 3 states. (Right) Each sub-track is assigned to the state for which it has the highest posterior probability. Reused from reference[7] (public domain). **(b)** Swarm charts of the maximum posterior probability for each pEMv2 state for the indicated datasets. **(c)** Ensemble mean squared displacement for ground truth (dashed lines) and QuantiTrack (solid lines) datasets for the indicated combinations of slow and fast states. Error bars = s.e.m. **(d)** Representative sub-tracks classified into the slow and fast states for the indicated mixtures. Scale bar = 1 μm.

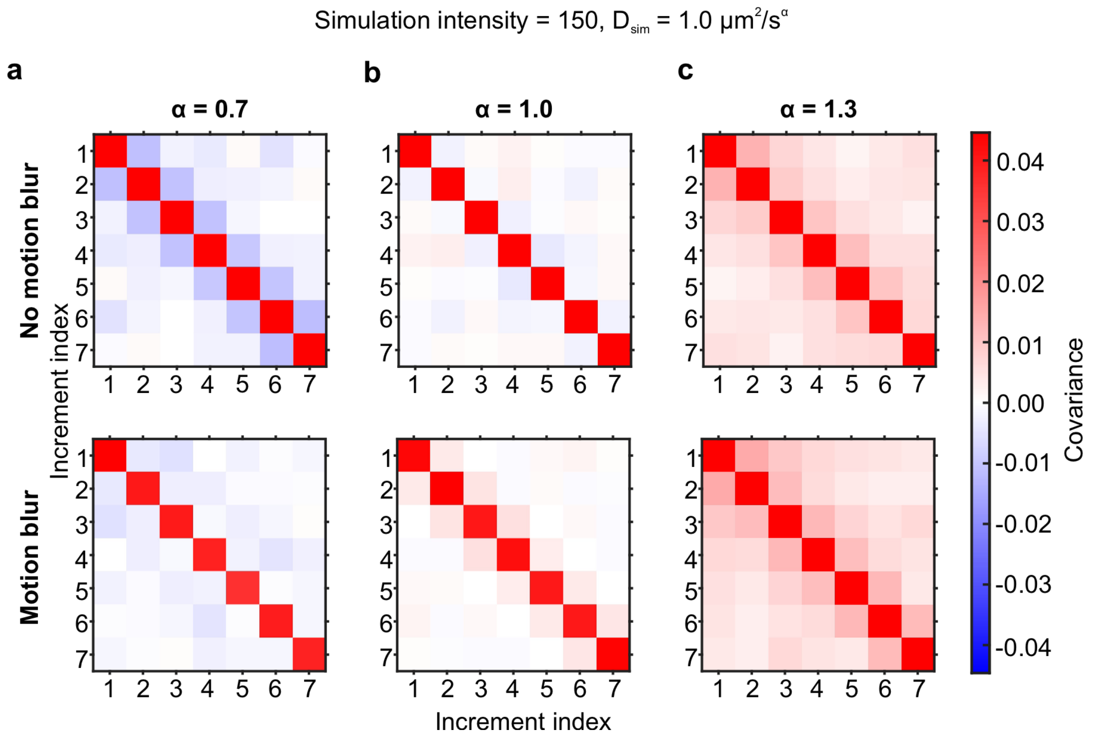


**Supplementary Figure 7. Covariance matrices for fractional Brownian motion.**
**(a–c)** Covariance matrix across seven step increments for (a) sub diffusion ($\alpha$ = 0.7), (b) normal diffusion ($\alpha$ = 1.0), and (c) super-diffusion ($\alpha$ = 1.3). (Top) Simulations without motion blur and (bottom) those with motion blur. $D_\alpha$ = 1.0 $\mu m^2/s^\alpha$.

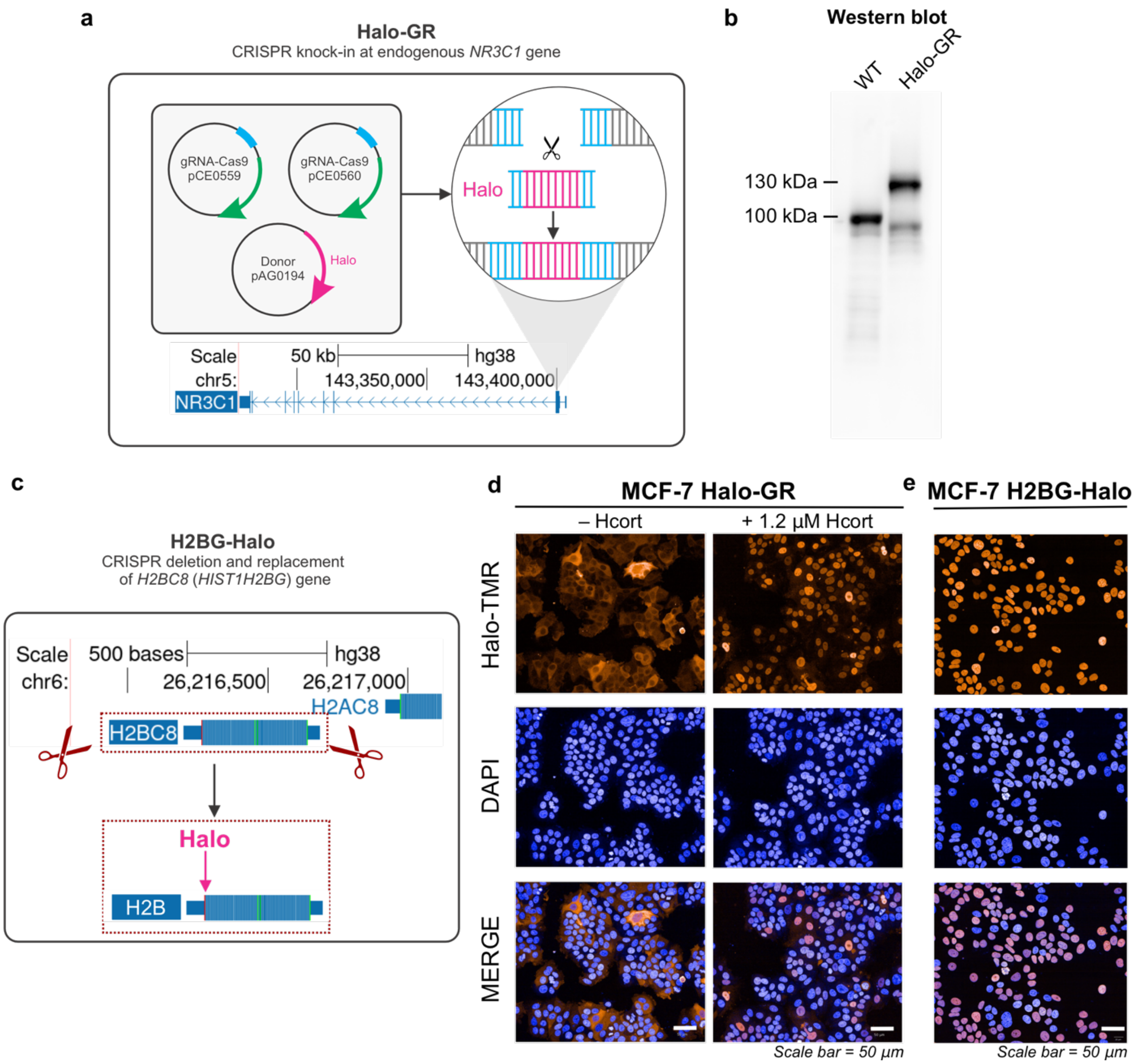


**Supplementary Figure 8. Generation and validation of Halo-GR and H2B-Halo MCF-7 cell lines.**
**(a)** To tag the endogenous GR in MCF-7 cells, the HaloTag cassette was inserted at the 5' end of the *NR3C1* gene (which encodes the glucocorticoid receptor [GR]) using CRISPR-Cas9. **(b)** Western blot for GR in the parental MCF-7 cells (WT) and clonal Halo-GR CRISPR knock-in cell line (Halo-GR). Expected sizes: 97 kDa (GR), 130 kDa (Halo-GR). **(c)** To generate the H2B-Halo cell line, the *H2BC8* gene was excised and replaced with an *H2B-HALO* gene using CRISPR-Cas9 and is expressed from the endogenous promoter. **(d)** Representative images of MCF-7 cells expressing Halo-GR. Halo-TMR staining (which covalently binds to the HaloTag) is shown in orange and DAPI staining is shown in blue. The column on the left shows the cells before hormone treatment, where GR is primarily cytoplasmic. The column on the right shows GR after treatment with 1.2 µM hydrocortisone (Hcort), which results in nuclear translocation of GR. **(e)** Representative images of H2B-Halo expressing MCF-7 cells. Halo-TMR staining is shown in orange while DAPI staining is shown in blue. Scale bars for panels d and e are 50 µm.

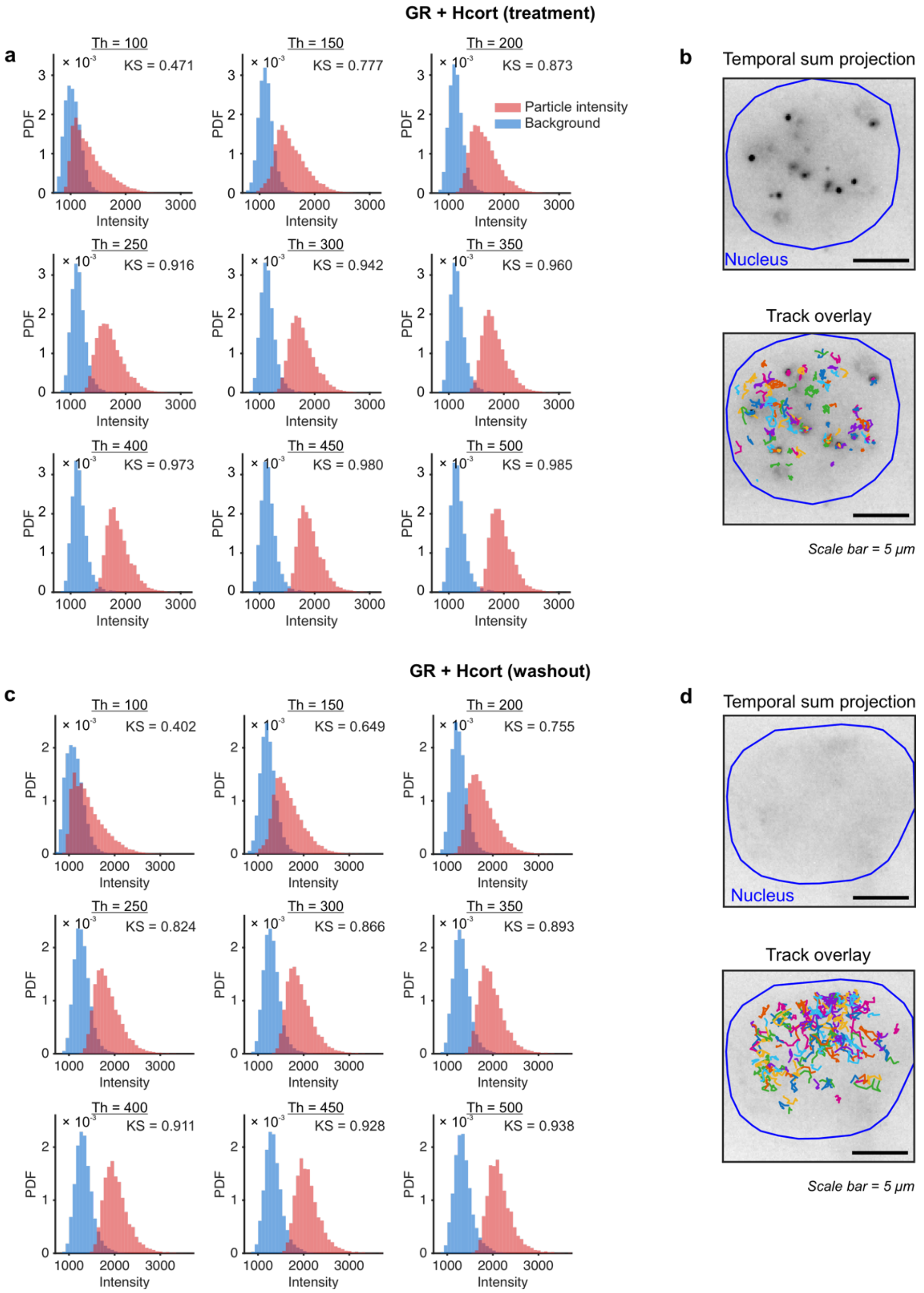


**Supplementary Figure 9. Particle signal and local background intensity distributions for fast GR SMT data.**

**(a)** Particle signal (red) and local background (blue) intensity distributions for GR + hydrocortisone (Hcort) treatment at indicated detection threshold values. Inset text shows the Kolmogorov-Smirnov (KS) statistic that measures the distance between the signal and background intensity distributions. **(b)** (Top) Temporal sum projection of a representative GR + Hcort treatment movie and (bottom) sum projection with the trajectories overlaid. Nucleus outline is shown in blue. Scale bar = 5 μm. **(c)** Particle signal (red) and local background (blue) intensity distributions for GR + hydrocortisone (Hcort) washout at indicated detection threshold values. (Inset) KS statistic. **(d)** (Top) Temporal sum projection of a representative GR + Hcort washout movie and (bottom) sum projection with the trajectories overlaid. Nucleus outline is shown in blue. Scale bar = 5 μm.

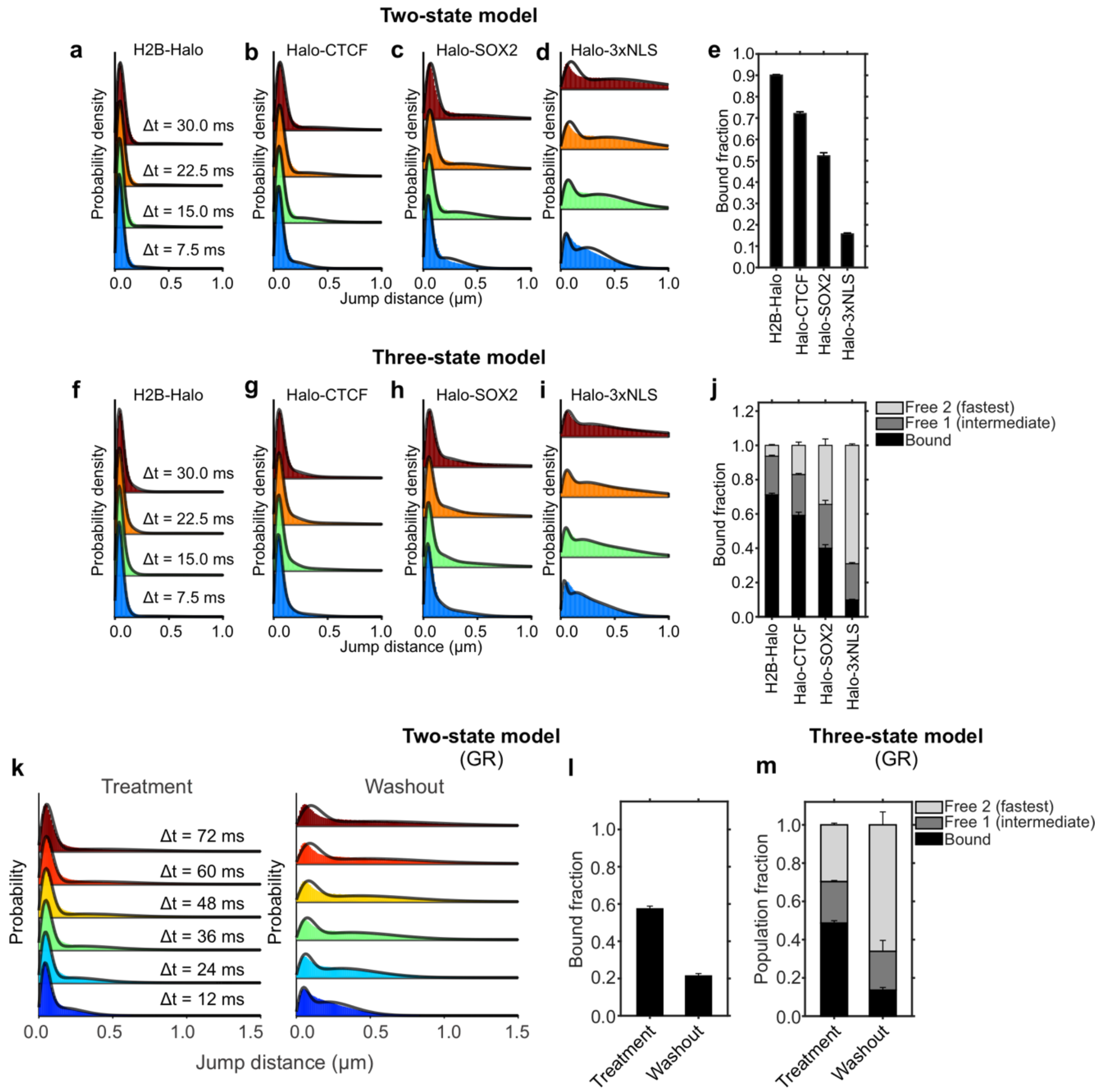


**Supplementary Figure 10. Spot-On models applied to nuclear proteins.**
**(a–d)** Jump distance distributions of (a) histone H2B-Halo, (b) Halo-CTCF, (c) Halo-SOX2, and (d) Halo-3xNLS with Spot-On two-state model fits (solid lines). **(e)** Measured bound fractions. **(f–i)** Jump distance distributions of (f) histone H2B-Halo, (g) Halo-CTCF, (h) Halo-SOX2, and (i) Halo-3xNLS with Spot-On three-state model fits (solid lines). **(j)** Measured population fractions. **(k)** Jump distance distributions and Spot-On two-state model fit (solid lines) for (left) GR + hydrocortisone (Hcort) treatment and (right) washout. **(l)** Bound fraction as measured by the two-state model. **(m)** Population fractions for the bound and two free states determined by Spot-On. Error bars in panels e, j, l, and m = bootstrapped standard deviation across 100 resamples.

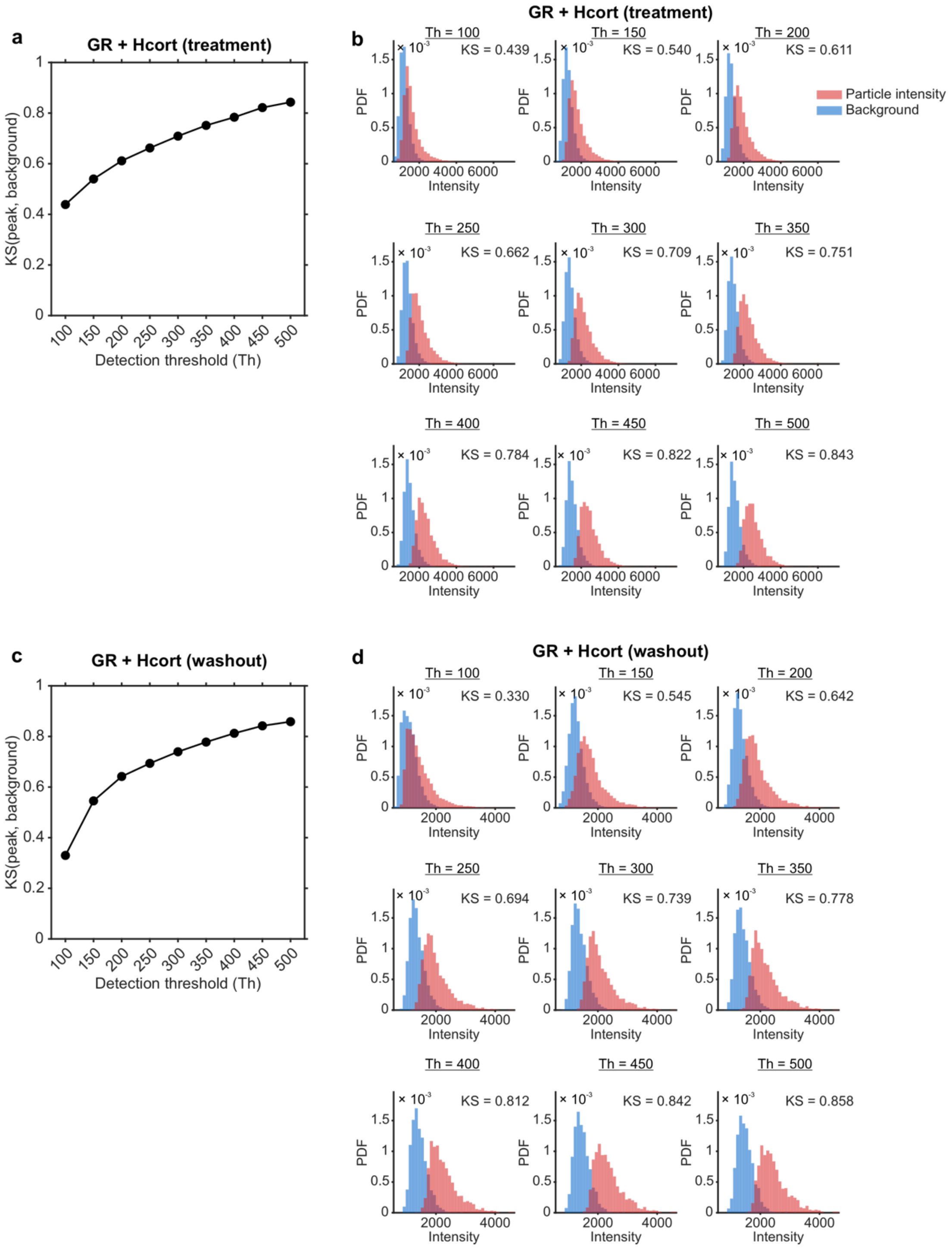


**Supplementary Figure 11. Signal and background intensity distributions for slow GR SMT data.** **(a)** Kolmogorov-Smirnov (KS) statistic quantifying the distance between the particle signal and background intensity distributions as a function of detection threshold (Th) for GR + Hcort treatment. **(b)** Particle signal (red) and local background (blue) intensity histograms at indicated detection thresholds for GR + Hcort treatment. **(c)** KS statistic as a function of Th for GR + Hcort washout. **(d)** Signal and background histograms at indicated Th for GR + Hcort washout.

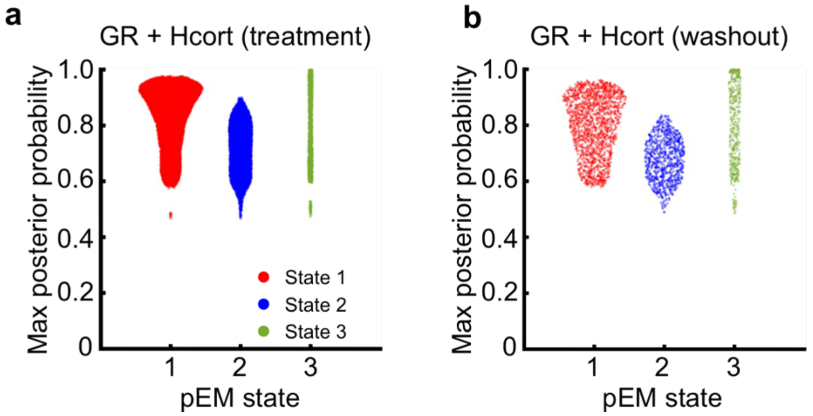


**Supplementary Figure 12. Perturbation expectation maximization v2 (pEMv2) analysis applied to the slow GR SMT data.**
**(a–b)** Swam charts of the maximum posterior probability for each pEMv2 state for GR under (a) Hcort treatment and (b) Hcort washout.

**Movie 1**
(2+1) D movie of GR + Hcort obtained with 12 ms exposures and continuous illumination.

**Movie 2**
Montage of simulated SMT movies of normal diffusion ($\alpha$ = 1, D = 1 $\mu m^2/s$) with motion blur and indicated signal intensity (Int). Scale bar = 5 $\mu$m.

**Movie 3**
Montage of simulated SMT movies of normal diffusion ($\alpha$ = 1, signal intensity = 150) with motion blur and diffusion coefficients (D) = {1.0, 2.5, 5.0, 7.5, 10.0, 12.5, 15.0, 20.0} $\mu m^2/s$. Scale bar = 5 $\mu$m.

**Movie 4**
Montage of simulated SMT movies of normal diffusion ($\alpha$ = 1, signal intensity = 150) without motion blur and diffusion coefficients (D) = {1.0, 2.5, 5.0, 7.5, 10.0, 12.5, 15.0, 20.0} $\mu m^2/s$. Scale bar = 5 $\mu$m.

**Movie 5**
Fast SMT movie of Halo-GR + Hcort treatment (left) and washout (right) overlaid with detected tracks (red). Only tracks longer than 4 frames are shown. Scale bar = 2 $\mu$m.

**Movie 6**
Slow SMT movie of H2B-Halo (left), Halo-GR + Hcort treatment (middle), Halo-GR + Hcort washout (right) overlaid with detected tracks (red). Only tracks longer than 4 frames are shown. Scale bar = 2 $\mu$m.
